# Comparative immuno-biology at clinical recognition of early multiple organ dysfunction syndrome in pediatric and adult patients using single-cell transcriptomics

**DOI:** 10.1101/2025.01.02.630924

**Authors:** Rama Shankar, Austin W. Goodyke, Shubham Koirala, Shreya Paithankar, Ruoqiao Chen, Nicholas L. Hartog, Dave Chesla, Surender Rajasekaran, Bin Chen

## Abstract

Globally, sepsis remains a major health issue, with Multiple Organ Dysfunction Syndrome (MODS) being a leading cause of mortality. MODS, a severe condition often seen in intensive care units, typically results from infections or trauma and involves complex pathophysiological processes requiring various clinical interventions. Although infections are the main triggers, the mechanisms driving MODS remain unclear. To investigate the transition of sepsis to MODS, we generated a single cell RNA sequencing dataset comprising 86,839 immune cells from pediatric sepsis patients at the clinical onset of MODS patients and age-matched controls, identifying 22 distinct cell types. A cluster of *S100* genes, located in the same genomic region, was highly expressed in neutrophils in MODS patients, demonstrating strong diagnostic potential across cohorts (AUC=0.94– 0.99) and potential as therapeutic targets. We found that many B and T cells showed heightened inflammation and increased apoptotic activity during early MODS. Additionally, specific transcription regulators and surface proteins associated with inflammation and *S100* regulations were uniquely expressed in MODS. Pseudotime analysis revealed distinct *S100* gene expression patterns between controls and MODS. Cell-cell interaction analysis highlighted dendritic cells as key mediators, enhancing communication between plasma cells and Vδ T cells while activating inflammatory and immunosuppressive pathways. We also analyzed 116,803 immune cells from adult MODS patients, revealing stronger immune dysregulation compared to pediatric MODS, including altered *S100* gene expression, and enhanced cell-cell interactions. These findings suggest that *S100* genes may serve as a marker for MODS. Furthermore, insights gained from adult MODS could improve our understanding of rare pediatric MODS and contribute to the development of better therapeutics for all MODS patients.

## Introduction

Multiple organ dysfunction syndrome (MODS) occurs in more than 25% of patients in pediatric intensive care units (PICUs) ^1^. It commonly arises in response to pro-inflammatory insults such as sepsis or trauma ^2,3^. In some sepsis patients, MODS results in a systemic pathologic state adversely affecting numerous organ systems simultaneously ^4^. Contemporary management of MODS is entirely supportive and focused on addressing the underlying disease process. That support could be organ specific such as dialysis or life preserving such as extracorporeal membrane oxygenation (ECMO) support ^5^. There is a financial burden as critically ill patients who develop MODS stay three times longer in the ICU and suffer higher mortality rates ^3,6^. It has also been observed that an early diagnosis of MODS would result in a decrease in severity and mortality ^7^. However, lack of a complete molecular understanding led to design various multiple organ failure (MOF) scoring systems to assess severity and facilitate risk stratification in critically ill patients ^8^. These scoring systems are tied into epidemiology of sepsis rather than MODS in specific. A deeper understanding of the molecular basis behind the transition from sepsis to MODS from multiple cohorts would be beneficial to designing better scoring systems and developing more effective therapies.

The development of MODS is widely attributed to uncontrolled immune system dysfunction, precipitated by the release of damage-associated molecular patterns (DAMPs) from extensive tissue damage ^9–11^. The degree of immune disruption that characterizes patients with MODS that required ECMO has been examined in a previous study ^12^, where we performed bulk RNA-seq data analysis on pediatric MODS patients at different time points. We observed that MODS patients exhibited elevated levels of neutrophils and monocytes, increased activation of NF-κB pathways and inflammatory responses, as well as heightened activity in cytokine and interferon-related pathways compared to controls ^12^. However, deeper insights into the mechanism of MODS initiation and more effective clinical interventions are needed. There remains a debate about whether initiation of MODS involves hyperinflammation, immunosuppression or some combination of both ^13^. In addition, the role of other immune cells during the MODS initiation and progression remains unknown. Circulatory immune cells are thought to be drivers of inflammation in MODS ^14^. Additionally, cellular and transcriptional differences between pediatric and adult MODS are also not well understood. Insights into the molecular characteristics of the immune system could be gained through advanced single-cell RNA sequencing (scRNA-seq).

In this study, we recruited five pediatric controls and five rare pediatric sepsis patients transitioning to MODS, matched for age and biological sex, and generated scRNA-seq data from the CD45+ cell population. Inspired by the copy number analysis in cancer, we identified a cluster of genes associated with MODS and evaluated their translational potential using multiple bulk transcriptomics datasets. We further investigated the inflammasome, immune cell activation, inferred surfaceome, transcriptional regulation, and key signaling pathways involved in MODS. Finally, we compared these findings with data from adult sepsis patients transitioning to MODS, highlighting the similarities and differences between pediatric and adult MODS.

## Results

### Immune cell landscape in MODS

We enrolled five pediatric MODS patients and five age- and sex-matched sedations (control) individuals (Table S1). For all MODS patients, a comprehensive record of 11 distinct clinical features was documented. Blood samples were collected from all ten individuals, from which CD45 positive cells were subsequently isolated and scRNA-seq data was generated (Fig. 1A). We adopted a modified protocol tailored for neutrophils to process the scRNA-seq data using CellRanger ^15^. Standard filtering criteria were applied to exclude cells with high mitochondrial genes, doublets, and empty droplets, resulting in a total of 86,839 cells from both control and MODS patients. Clustering of cells based on patient groups indicated minimal batch effects (Fig. 1B).

**Fig. 1.**
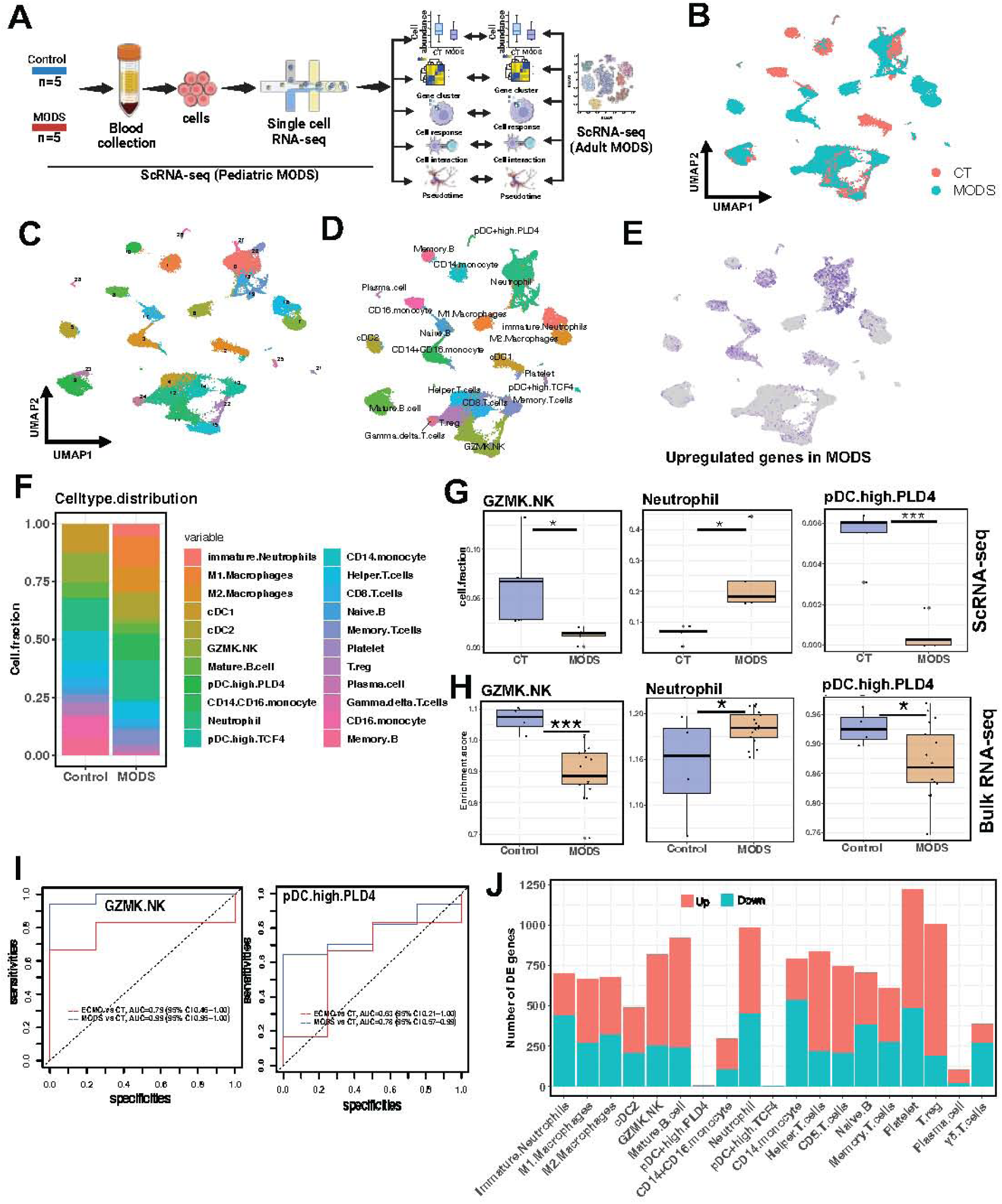
scRNA-seq in pediatric MODS. (A) Schematic of the study. (B-C) UMAP plot displaying cell clusters colored by disease groups (B) and Seurat clustering (C). (D) UMAP plot illustrating clustering of identified cell types. (E) Enrichment of upregulated genes obtained from bulk RNA139 seq by comparing MODS with control samples from our previous study (Shankar et al., 2020). (F) Cell fraction distribution in control and MODS. (G) Cell types exhibiting differences in abundance between MODS and control based on scRNA-seq data. Each dot represents one sample. The black line indicates the median value, and the vertical line represents the standard deviation. (H) Cell type enrichment estimated from the deconvolution of bulk RNA-seq data. Gene markers for all three cell types were used to compute enrichment in bulk RNA-seq data. (I) Receiver operating characteristic (ROC) curves for the classification of MODS and control samples as well as MODS patients with ECMO support (ECMO) and MODS patients using the enrichment scores of GZMK+NK cells and pDC+highPLD4 cells. Area Under the ROC curve (AUC) was computed to quantify performance. (J) Number of DE genes between MODS and control in each cell type. *** 1e-05 < p-value ≤ 0.001 and * 0.01< p-value ≤ 0.05.

Further clustering analysis revealed 27 clusters (Fig. 1C), corresponding to 22 distinct cell types (Fig. 1D). The majority of cell types belonged to seven major classes: T lymphocytes (CD8 T cells, memory T cells, GZMK.NK cells, Vδ T cells, helper T cells, and T regulatory cells), B cells (mature B cells, memory B cells, and naïve B cells), plasma cells, monocytes (CD16+CD14 monocytes, CD14 monocytes, CD16 monocytes, M1 macrophages, and M2 Macrophages), dendritic cells (cDC1, cDC2, pDC+high TGF4, and pDC+high PLD4), neutrophils (mature and immature), and platelets (Fig. 1D).

Previous bulk RNA-seq data showed upregulated genes associated with various immune related processes in MODS ^12^. Therefore, we selected these upregulated genes and explored their enrichment in our dataset. Neutrophils and monocytes were identified as major contributors to these genes in MODS (Fig. 1E), corroborating with the deconvolution analysis of bulk RNA-seq ^12^. Cell fraction analysis revealed altered cell distributions in MODS compared to control (Fig. 1F and Figure S1), with a decrease in GZMK.NK cells and pDC+high PLD4 cells, and an increase in neutrophils (Fig. 1G). These observations were further validated using FACS data (Figure S1). Deconvolution of an independent bulk RNA-seq dataset from our previous study ^12^ using gene markers of these cells revealed a similar pattern (Fig. 1H). Notably, the enrichment of GZMK.NK cells and pDC+high PLD4 cells significantly differentiated control and MODS patients, as indicated by the area under the curve (AUC) values (0.99 for GZMK.NK and 0.78 for pDC+high PLD4) (Fig.1I). This suggested that the abundances of GZMK.NK and pDC+high PLD4 may be further explored as biomarker candidates for the diagnosis of pediatric ICU patients progressing towards MODS. Additionally, we examined the number of DE genes in each cell type by comparing cells from MODS with control. Platelets, T regulatory cells (Tregs), and neutrophils exhibited the most extensive transcriptional changes, whereas pDCs, CD14/CD16+ monocytes, and plasma cells showed the least transcriptional alterations in MODS (Fig.1J).

### Cluster of S100 genes overexpressed in MODS

Although many cell types exhibited significant transcriptional changes in MODS, the heterogeneity of MODS prompted us to explore transcriptional changes across cells in individual patients. Interestingly, we found the InferCNV package ^16^, originally developed to infer copy number variations (CNVs) from gene expression in tumor cells, could reveal distinct clusters shared by individual MODS patients. InferCNV estimates CNVs by analyzing consistent over- or under-expression of genes in a genomic region relative to a reference. Using cells from control samples as the reference, we observed that several regions on chromosome 1 clustered together in neutrophils from MODS patients, with these regions consistently upregulated (Fig. 2A for one patient). This pattern was consistent across all MODS patients (Figure S2). Further annotation of these genomic locations revealed that 13 genes (*CTSS, C1orf56, S100A4, S100A6, S100A8, S100A9, S100A10, S100A11, S100A12, SH2D2A, FCRL1, FCRL3,* and *MNDA*) were consistently upregulated in neutrophils in every MODS patient (Fig. 2B and Figure S3).

**Fig. 2.**
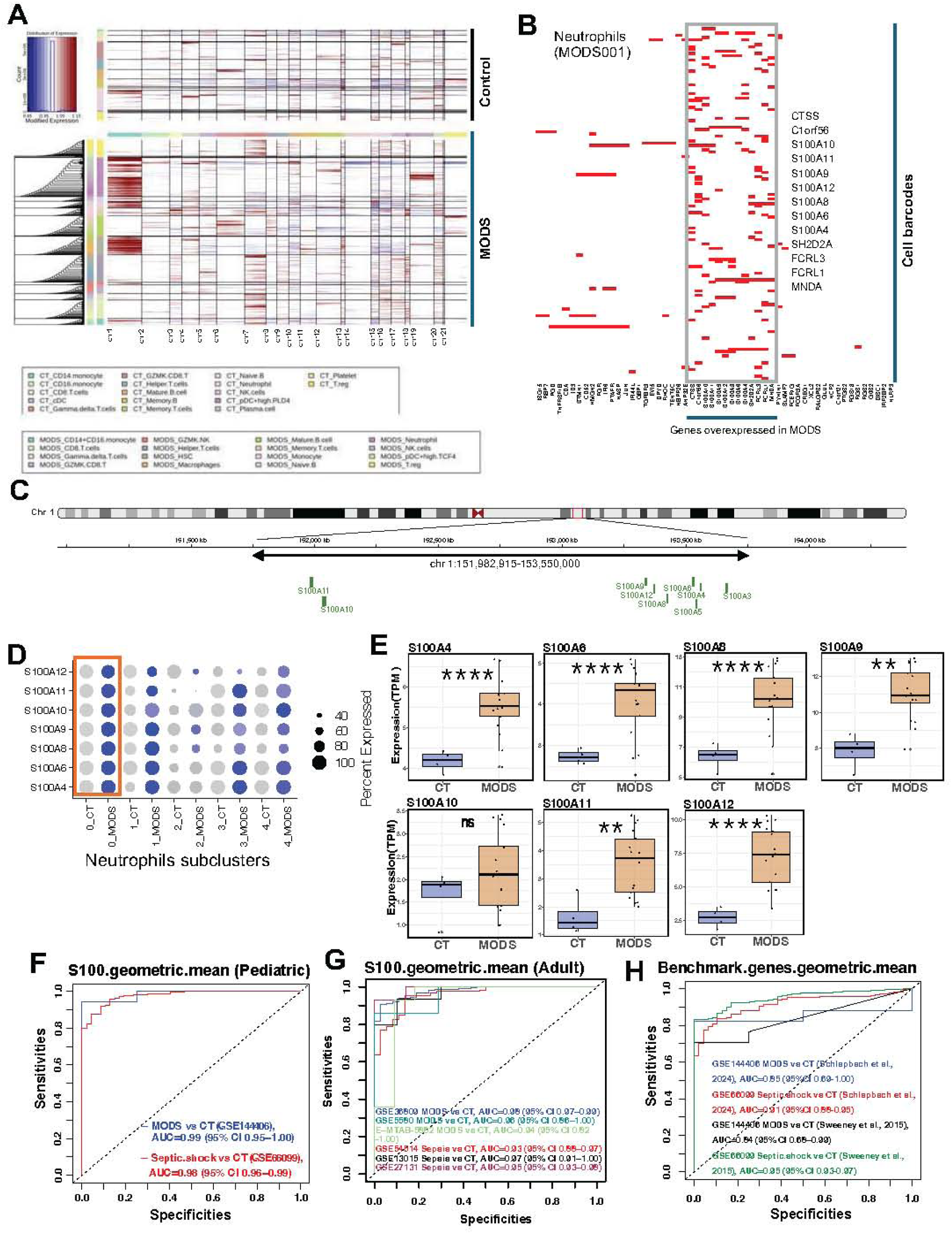
S100 gene cluster overexpression in pediatric MODS. (A) InferCNV results of MODS001 imply enhanced co-expressed cluster genes in MODS within neutrophils. The co-expressed region was consistently observed in neutrophils across all MODS patients (Figure S2). (B) Highly expressed genes in neutrophils from a single MODS patient (MODS001). These genes exhibited consistent overexpression across all MODS patients (Figure S2-S3). (C) UCSC genome browser screenshot depicting S100 genes located on chr 1:151,982,915-153,550,000. The blue bars represent S100 genes. (D) Expression of S100 genes in different subclusters of neutrophils. The expression pattern of these genes in each cell type is depicted in Figure S4A. Grey color indicates no expression, while blue color represents high expression of S100 genes. The size of the circles indicates the percentage of cells expressing individual genes in each cluster. (E) Expression of S100 genes in bulk RNA-seq data from a different study. Except for S100A10, all genes were significantly upregulated in MODS. (F-G) ROC curves for differentiating MODS patients in pediatric (F) and adult (G) using the geometric mean of the expression of S100 genes. (H) ROC curves for differentiating pediatric control and MODS samples using benchmark genes from two different studies. AUC was computed to quantify performance. **** 1e-16< p-value ≤ 1e-05, *** 1e-5 < p-value ≤ 0.0001, ** 0.0001 < p-value ≤ 0.01, * 0.01< p-value ≤ 0.05.

Interestingly, only the S100 genes were located in the same region (Fig. 2C) and were upregulated in neutrophils (mature and immature), specifically in cluster0 of mature neutrophils (after re-clustering the neutrophils) (Fig. 2D and Figure S4A).

Further analysis using bulk RNA-seq data revealed that six out of seven S100 genes were significantly upregulated in MODS (Fig. 2E). Since, S100 genes are classified as DAMPs ^17,18^, we also investigated the expression of other DAMPs such as histone genes, heat shock proteins (HSPs), *HMGB1*, and *HMGN1*. However, no significant differences were observed in the expression of these genes (Figure S4B), indicating that *S100* genes are the primary DAMP-associated genes at initiation of MODS.

The expression of *S100* genes increased in trauma patients from the time of admission to 72 hours of hospitalization progressing toward MODS, while this pattern was not observed in control or ICU patients who did not develop MODS in another cohort (Figure S4C). Furthermore, the geometric mean expression of these S100 genes could significantly differentiate MODS patients from control (AUC=0.99 in GSE144406 (pediatric patients), AUC=0.98 in GSE36809, AUC=0.96 in GSE5580, and AUC=0.94 in E-MTAB-5882) as well as septic shock patients from control (AUC=0.98 in GSE66099 (pediatric patients), AUC=0.93 in GSE54514, AUC=0.97 in GSE13015, and AUC=0.95 in GSE27131)) across multiple datasets (Fig. 2F-G). S100 genes outperformed multiple benchmark gene sets proposed previously ^19,20^ in distinguishing control and MODS patients (AUC varies from 0.84-0.85 for control vs MODS in GSE144406 and AUC varies from 0.91-0.95 for control vs septic shock in GSE66099), suggesting that *S100* genes may serve as a potential biomarker for the early diagnosis of MODS patients.

### Inflammasome in MODS

Sepsis is fundamentally an inflammatory disease driven by the host immune response. Since *S100* genes are well-known DAMP genes linked to inflammation, we investigated the cells associated with inflammasome activity. We observed increased inflammasome activity in CD8+ T cells, GZMK+ NK cells, helper T cells, memory T cells, mature B cells, and pDC+highPLD4 cells in MODS (Fig. 3A). Conversely, naïve B cells, Treg cells, and neutrophils exhibited reduced inflammasome activity (Figure S5A). Alternatively, CD8 T cells, neutrophils, GZMK+ NK cells, and Treg cells showed greater immune activation in MODS (Fig. 3B), while other cell types displayed reduced activation (Figure S5B), highlighting inflammation as a pivotal event in the initiation of MODS.

**Fig. 3.**
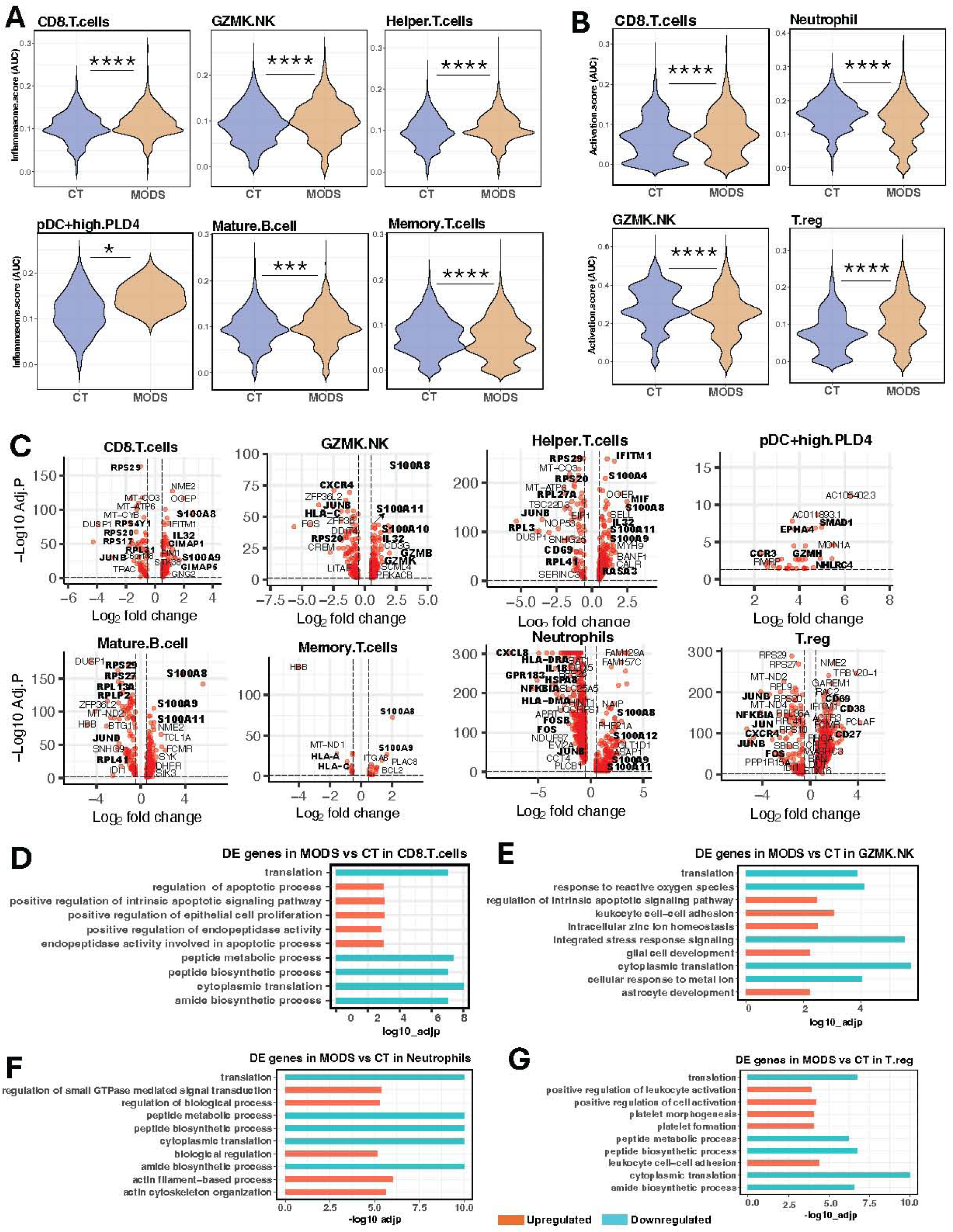
Role of immune cells in inflammation and immune activity. Inflammasome (A) and immune cell activation (B) activities in immune cells from control and MODS groups. (C) Volcano plot highlighting differentially expressed genes across various cell types in MODS. (D-G) Enriched biological processes identified by upregulated (red) and downregulated (blue) genes in CD8+ T cells (D), GZMK+ NK cells (E), neutrophils (F), and Treg cells (G). **** 1e-16< p-value ≤ 1e-05, *** 1e-5 < p-value ≤ 0.0001, * 0.01< p-value ≤ 0.05.

Differential gene expression analysis by comparing cells from MODS with control revealed that many of these cells expressed *S100* genes (*S100A8, S100A9, S100A10, S100A12*) (Fig. 3C). Additionally, CD8+ T cells, GZMK+ NK cells, helper T cells, mature B cells, memory T cells, and pDC+highPLD4 cells exhibited higher expression of inflammasome-related genes such as *JUN, JUNB, MIF,* and *IL32* in MODS.

In contrast, neutrophils and Treg cells showed downregulation of these genes. Notably, Treg cells displayed upregulation of *CD27*, *CD38*, and *CD69,* which are markers associated with T cell activation and differentiation. Neutrophils showed reduced expression of HLA genes (*HLA-DRA* and *HLA-DMA*), *CXCL8*, and *HSPA8* (Fig. 3C).

Further, pathway analysis of DE genes in each cell type indicated widespread downregulation of protein and peptide synthesis across most of cell types (Fig. 3D-G), potentially as an adaptive mechanism to limit immune hyperactivity and pathogen infection ^21^. However, CD8+ T cells, helper T cells, naïve B cells, and GZMK+ NK cells demonstrated upregulated apoptotic processes (Fig. 3D-G, Figure S5C-G). Treg cells showed increased leukocyte activation and platelet formation, while neutrophils exhibited upregulation of GTPase-mediated signaling (Fig. 3F). Mature B cells displayed enhanced inflammatory responses, whereas naïve B cells showed downregulation of antigen presentation *via* MHC class II genes (Figure S5D-F). Interestingly, all immune cell types exhibited increased apoptotic activity in MODS (Figure S6), suggesting a dual mechanism: reducing protein synthesis to mitigate hyperactivity while promoting apoptosis to eliminate damaged cells and facilitate phagocytosis.

Given the observed changes in MHC expression and signaling genes, we further examined cell surface protein expression in each cell types. CD8 T cells, memory T cells, and helper T cells showed upregulation of IL2RB, NCR3, PTP4A2, PVRIG, CD247, and KLRK1, while CD4 and CD325 were downregulated (Figure S7). Neutrophils exhibited reduced expression of HLA-DRB5, HLA-DQA2, IgG, and CAM2D, while GZMK+ NK cells displayed downregulation of IgG alone (Figure S7).

### Alternation of cell stages and regulatory network in MODS

Given the heightened apoptotic activity observed in MODS, we investigated cellular turnover using pseudotime analysis. Neutrophils and immature neutrophils exhibited the highest expression levels of stemness-associated genes. Using immature neutrophils as a reference, we observed that in control samples, cellular differentiation occurred simultaneously, with pseudotime trajectories diverging into multiple branches, indicating the maturation of various immune cell lineages (Fig. 4A-B). In contrast, in MODS, pseudotime trajectories predominantly progressed towards B and T lymphocytes, specifically memory T cells, CD8 T cells, naïve B cells, and helper T cells at very early pseudotime (Fig. 4A-B), which are critical components of the adaptive immune response essential to eliminate infected cells and pathogens during secondary infections.

**Fig. 4.**
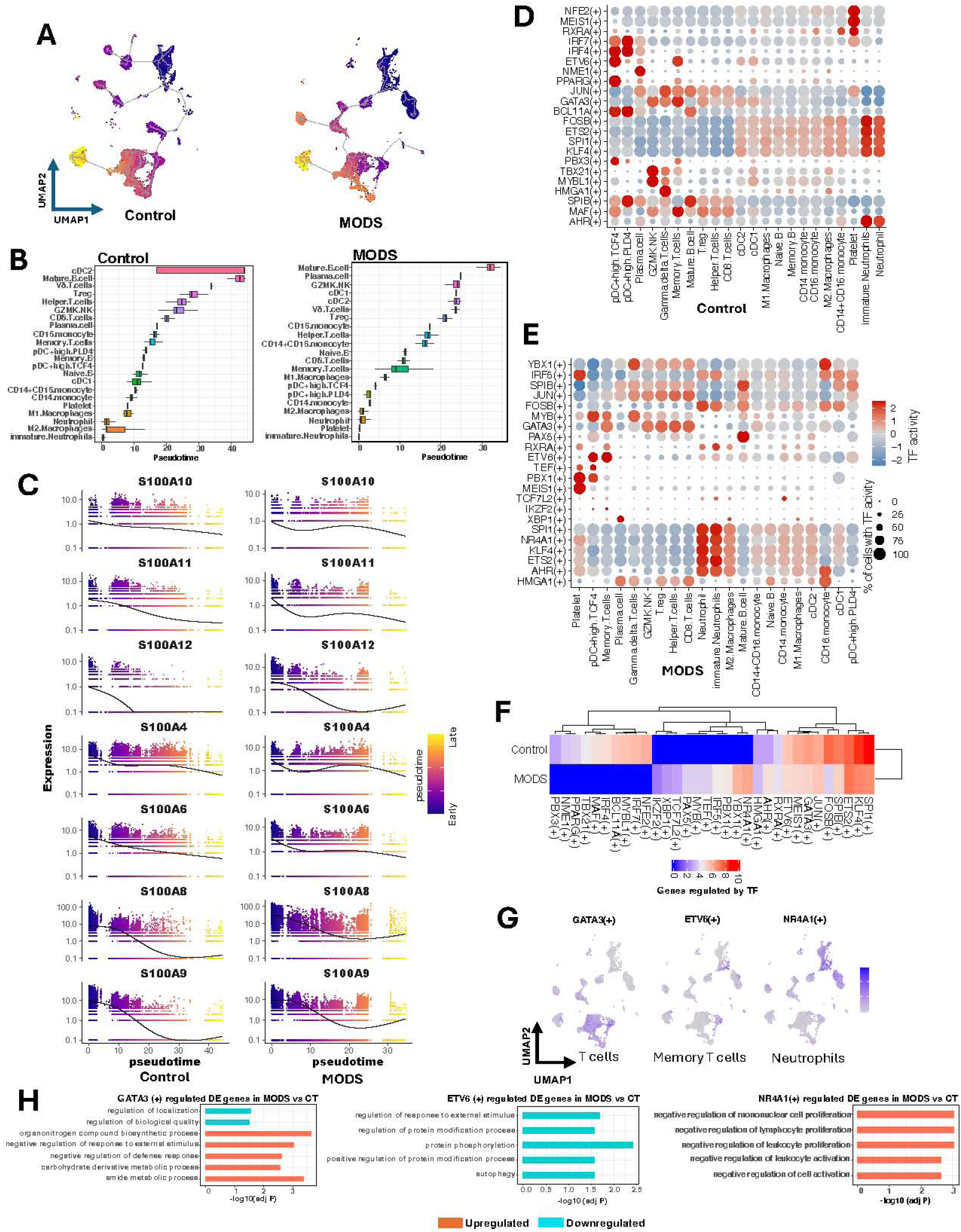
Pseudotime and active regulons in pediatric MODS. (A) Pseudotime analysis of cells in control and MODS, with blue indicating the pseudotime starting point and yellow marking the end stage. (B) Boxplot showing the median pseudotime of each cell type. (C) Expression changes of S100 genes in control (left) and MODS (right). (D-E) Dot plots showing the activity of active regulons in control (D) and MODS (E) samples. The size of each circle represents the percentage of cells with transcription factor (TF) activity, with blue indicating decreased TF activity and red indicating increased TF activity across different cell types. (F) Heatmaps displaying the number of regulating genes in control and MODS, where blue represents fewer or no regulating genes and red indicates a higher number of regulating genes. (G) Feature plots illustrating TF activity specific to T cells GATA3(+), memory T cells ETV6(+), and neutrophils NR4A1(+) in MODS. Blue indicates higher TF activity, while gray represents little or no TF activity. (H) Enriched biological processes associated with genes regulated in MODS (red) and in control (blue) by GATA3(+) (left), ETV6(+) (middle), and NR4A1(+) (right) regulons.

We further examined the variations in expression of *S100* gene cluster along pseudotime. In controls, *S100* genes were specifically expressed in neutrophils, with expression levels declining as pseudotime advanced (Fig. 4C). In MODS, *S100* gene expression followed a biphasic pattern: showing initial downregulation followed by reactivation at later pseudotime points. Among the *S100* genes in MODS, similar trends were noted for *S100A8, S100A9*, and *S100A12,* while *S100A4, S100A6, S100A10*, and *S100A11* displayed distinct expression trajectories (Fig. 4C) suggesting potential intra and inter interactions.

To elucidate the regulatory mechanisms underlying these genes and other differentially expressed genes, we inferred transcription factor (TF) activity in control and MODS samples, identifying 22 regulons with differential activity (Fig. 4D-E). In controls, pDCs demonstrated elevated activity of IRF4(+), IRF7(+), ETV6(+), PPARG(+), BCL11A(+), PBX3(+), and SPIB(+), which are critical for immune cell development, differentiation, and responses to inflammation (Fig. 4D). In MODS, neutrophils and immature neutrophils exhibited increased activity of SPI1(+), NR4A1(+), KLF4(+), ETS2(+), AHR(+), RXRA(+), FOSB(+), and IRF5(+) (Fig. 4E). Among these, SPI1(+), ETS2(+), FOSB(+), and IRF5(+) are potent activators of pro-inflammatory pathways ^22,23^, whereas NR4A1(+), KLF4(+), AHR(+), and RXRA(+) play regulatory or anti-inflammatory roles ^24,25^, underscoring the dual role of neutrophils in modulating inflammation during MODS onset. Comparison of regulons between control and MODS revealed that NR4A1(+), YBX1(+), PBX1(+), IRF5(+), TEF(+), MYB(+), PAX5(+), TCF7L2(+), XBP1(+), and IKZF2(+) were specifically active in MODS (Fig. 4F). These regulons are known to play critical roles in modulating inflammation by regulating cytokines and influencing the activity of other immune cells.

Further investigation into key TFs revealed that GATA3(+) and ETV6(+), predominantly expressed in T cells in MODS (Fig. 4G), suppressed genes involved in response to external stimuli and defense mechanisms, suggesting restricted T cell effector functions (Fig. 4H). In contrast, TFs expressed in neutrophils, including NR4A1(+), SPI1(+), KLF4(+), ETS2(+), and AHR(+) (Fig. 4G and Figure S8A), regulated pathways associated with IL-17 signaling, TOR signaling, protein synthesis, and leukocyte activation. Additionally, these regulons are known for their regulation of *S100* genes and inflammation. These pathways underscore their role in modulating immune responses and maintaining homeostasis during MODS initiation (Fig. 4H and Figure S8B-E). Collectively, these findings suggest that both neutrophils and lymphocytes actively balance immune hyperactivity through distinct regulatory mechanisms, contributing to immune homeostasis during the early stages of MODS.

### Cell-cell communication in MODS

Cell-cell communication (CCC) analysis among all the cell types revealed a reduction in overall communication in MODS (Fig. 5A). Specifically, interactions among immature neutrophils, neutrophils, and mature B cells were diminished, whereas interactions of pDC+highPLD4 with plasma cells with Vδ T cells were increased (Fig. 5B and Figure S9A). In control, mature B cells were the primary source of interactions, whereas in MODS, pDC+highPLD4 cells emerged as the major source of signaling (Figure S9B). The interaction of pDC+highPLD4 with plasma cells and Vδ T cells facilitated the release of signaling molecules including CCL, VISFATIN, CD40, PARs, APRIL, BTLA, and RESISTIN, which were specific to MODS samples (Fig. 5C and Figure S9C).

**Fig. 5.**
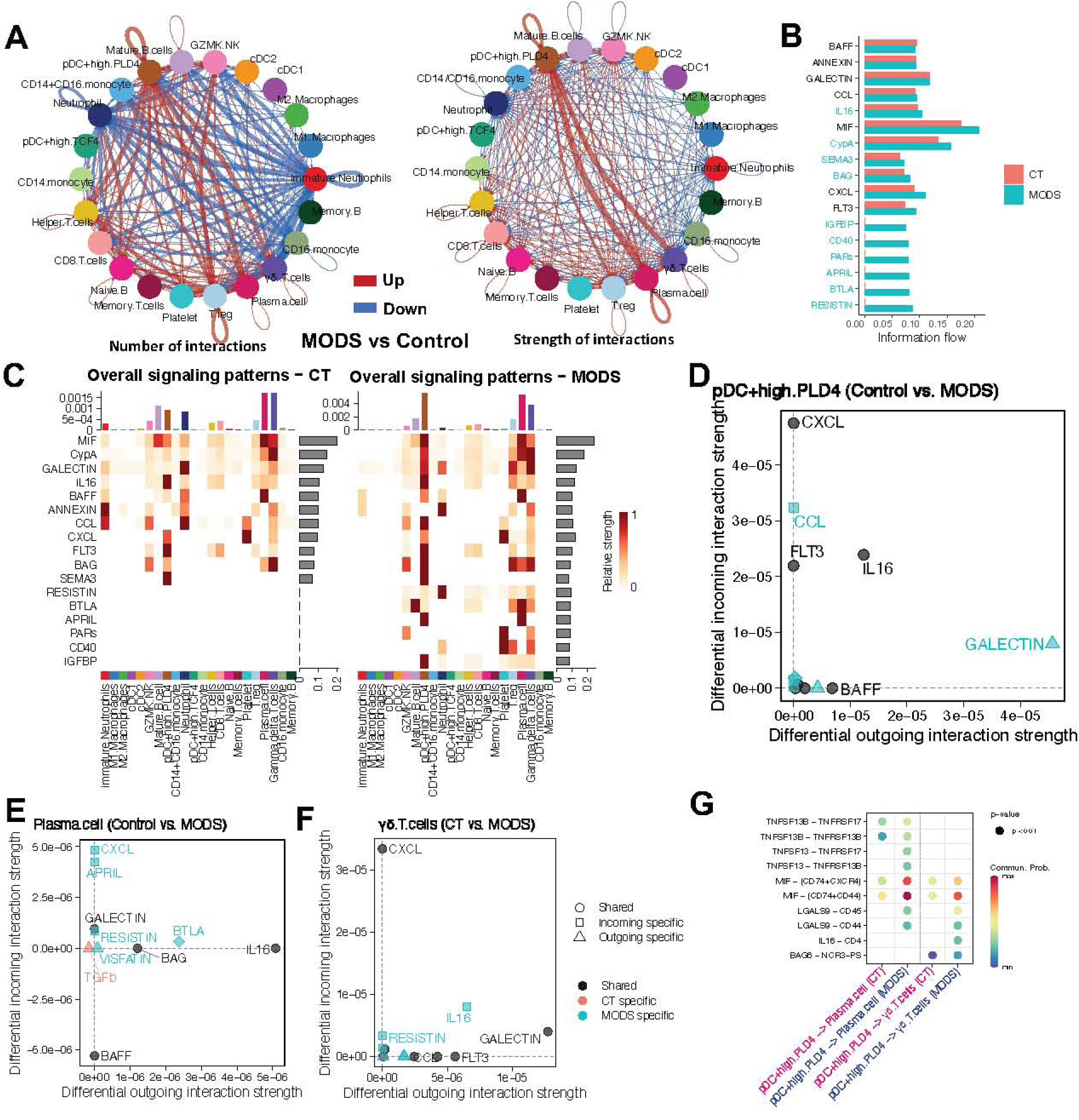
Cell-cell communications in pediatric MODS. (A) Differential cell-cell communications between MODS and control samples. The left panel of the cell-cell interaction network in (A) displays the number of interactions between two cell types, with red indicating an increase and blue indicating a decrease in cell interactions in MODS. The thickness of internodes represents the potential number of interactions (expressed ligand-receptor). The right panel of the network in (A) shows the interaction strength, with the thickness of the internode representing the strength of interactions. The strength of interactions was computed by the significantly co-expression of ligand and receptor pairs (log_2_ fold change ≥ 0.5 and p-value ≤ 0.05). (B) Signaling molecules in cell signaling. (C) Overall strength of signaling molecules in individual cell types. (D-F) Signaling molecules associated with signaling in pDC+highPLD4 cells (D), plasma cells (E), and γδ T cells (F). Molecules in blue are specific to MODS, molecules in grey are common to both control and MODS, whereas molecules in red are specific to control. (G) Expression of ligand receptor pairs associated with signaling in pDC+highPLD4 cells, plasma cells, and γδ T cells. Red represents higher probability and blue indicates lower probability.

Comparison of pDC+highPLD4 cells in MODS vs control revealed that CCL and GALECTIN were expressed specifically in MODS, while CXCL, FLT3, IL16, and BAFF were present in both (Fig. 5D). In MODS, plasma cells expressed signaling molecules including CXCL, APRIL, RESISTIN, VISFATIN, and BTLA (Fig. 5E), whereas Vδ T cells in MODS expressed IL16 and RESISTIN (Fig. 5F), indicating activation of specific signaling pathways in these cells. Further analysis of ligand-receptor pairs revealed that TNFS13-TNFRSF17, TNFS13-TNFRSF13B, LGALS9-CD45, and LGALS9-CD44 pairs mediated interactions between pDC+highPLD4 with plasma cells in MODS, whereas LGALS9-CD45, LGALS9-CD44, and IL16-CD4 pairs were associated with the interactions between pDC+highPLD4 and Vδ T cells in MODS (Fig. 5G).

### Immune dysregulation during the initiation of MODS in adult patients

To gain deeper insights into MODS, we analyzed a previously published scRNA-seq dataset ^26^ that included data from healthy adult controls and six disease conditions (details in methods) (Fig. 6A). We processed the dataset with Seurat, obtaining 116,803 high-quality cells with minimal batch effects (Fig. 6A). Clustering analysis identified 30 distinct cell clusters (Fig. 6B), which were subsequently annotated into 20 distinct cell types using Seurat’s label transfer approach (Fig. 6B). Further, we focused on the cells from control and ICU patients at the onset of MODS (Int-URO), as they shared a similar etiology with the studied pediatric MODS.

**Fig. 6.**
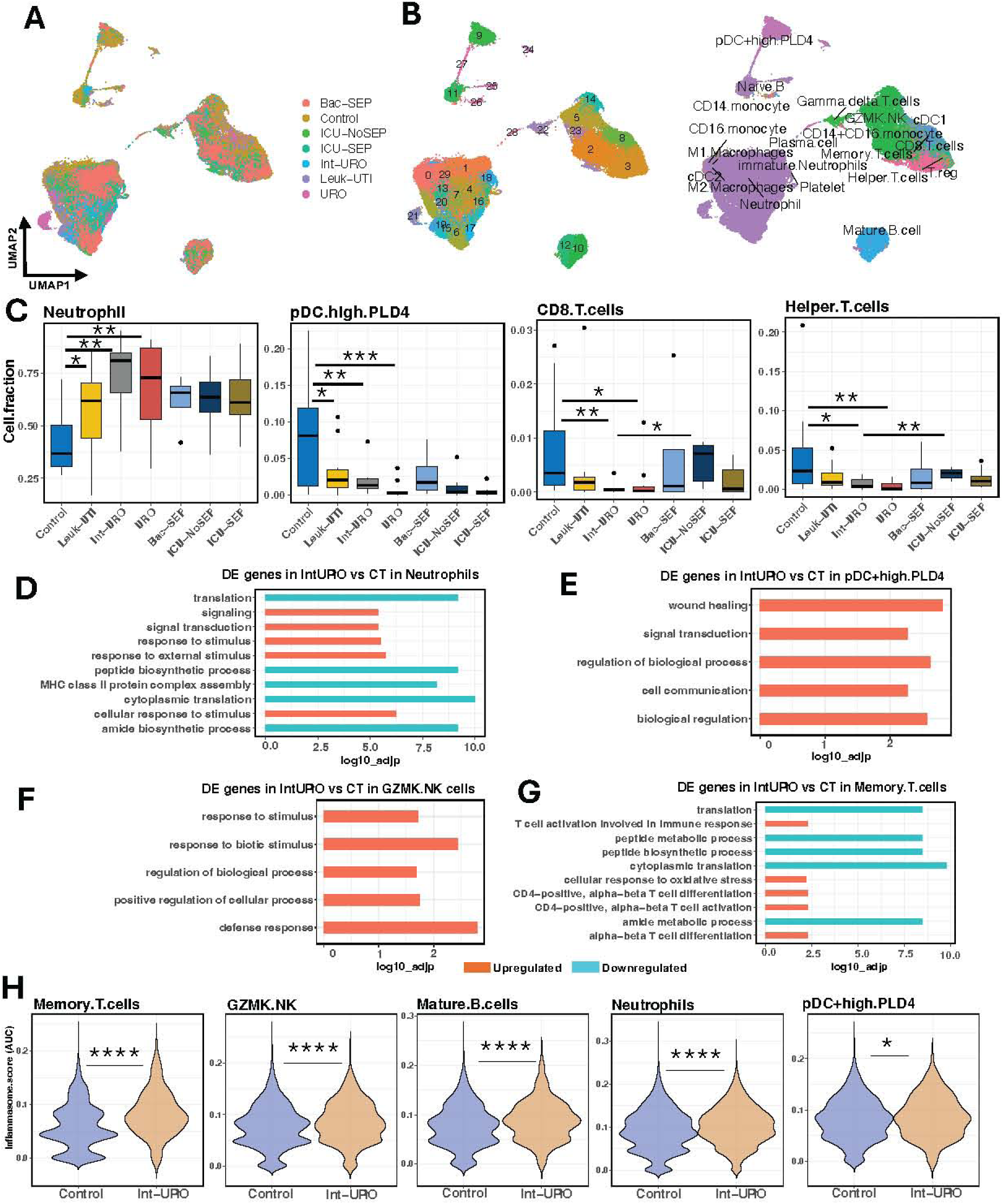
scRNA-seq in adult MODS. (A) UMAP plot displaying the clustering of cells, colored by sample type. (B) Cell clusters (left panel) and annotated cell types (right panel) in adult scRNA372 seq data. Cell types were annotated using the Seurat label transfer approach, with pediatric scRNA-seq data as the reference. Cells are colored by their assigned cell types. (C) Distribution of cell type fractions in control, Leuk-UTI, Int-URO, URO, Bac-SEP, ICU-NoSEP, and ICU-SEP, focusing on neutrophils, pDC+high PLD4, CD8+ T cells, and helper T cells. The distribution of other cell types is shown in Figure S10A. Significant differences are indicated with asterisks. (D-G) Enriched biological processes identified by upregulated (red) and downregulated (blue) genes, comparing Int-URO (MODS) with control in neutrophils (D), pDC+high PLD4 (E), GZMK+ NK cells (F), and memory T cells (G). Additional enriched processes are presented in Figure S10B. (H) Inflammasome activity in control and Int-URO samples for memory T cells, GZMK+ NK cells, mature B cells, neutrophils, and pDC+high PLD4. Except for memory B cells, all other cell types show decreased inflammasome activity in Int-URO. **** 1e-16< p-value ≤ 1e-05, *** 1e-5 < p-value ≤ 0.0001, * 0.01< p-value ≤ 0.05.

Cell proportion comparison between control and Int-URO revealed that neutrophils and pDC+high PLD4 cells in adult MODS exhibited a pattern similar to that observed in pediatric MODS (Fig. 6C). However, in contrast to pediatric MODS, no significant difference was observed in the abundance of GZMK+ NK cells between control and MODS in adult patients. Interestingly, cDC1, immature neutrophils, M2 macrophages, CD8+ T cells, helper T cells, and memory T cells showed reduced abundance in adult MODS patients (Fig. 6C and Figure S10A).

Comparative analysis of cells from control and MODS samples revealed that neutrophils exhibited downregulation of genes associated with protein synthesis and upregulation of genes related to signaling and responses to external stimuli (Fig. 6D). Similar trends were observed in the analysis of the DE genes in pDC+high PLD4, GZMK+ NK cells, cDC1, and mature B cells, where most of the upregulated genes were found to be associated with defense responses (Fig. 6E-F and Figure S10B). Memory T cells displayed the downregulation of genes involved in protein synthesis and upregulation of genes associated with T cell activation (Fig. 6G), indicating an adaptive response to eliminate infected cells and maintain immune homeostasis during MODS initiation. These findings were consistent with observations made in pediatric MODS.

Given the central role of inflammation in pediatric MODS, we investigated the inflammatory activity of immune cells in adult MODS. Surprisingly, except for memory T cells, no significant increase in inflammatory activity was observed in other immune cell types (Fig. 6H). However, neutrophils, GZMK+ NK cells, Vδ T cells, and Tregs exhibited enhanced immune activity (Fig. 10C). Collectively, these findings suggest that inflammation may not be the primary driver of MODS initiation in adult patients.

### S100 genes activity and cell-cell communications in adult MODS

Given the activation of *S100* gene cluster in pediatric MODS, we also investigated their expression in adult MODS. Similar to pediatric MODS, *S100* genes were predominantly expressed in neutrophils and immature neutrophils (Fig. 7A). When analyzing the expression of *S100* genes during disease progression, control samples exhibited minimal expression. However, as sepsis advanced towards MODS initiation (Int-URO), *S100* gene expression increased monotonically, peaking at the onset of MODS (Int-URO samples) and further declining in established MODS patients (URO) (Fig. 7B).

**Fig. 7.**
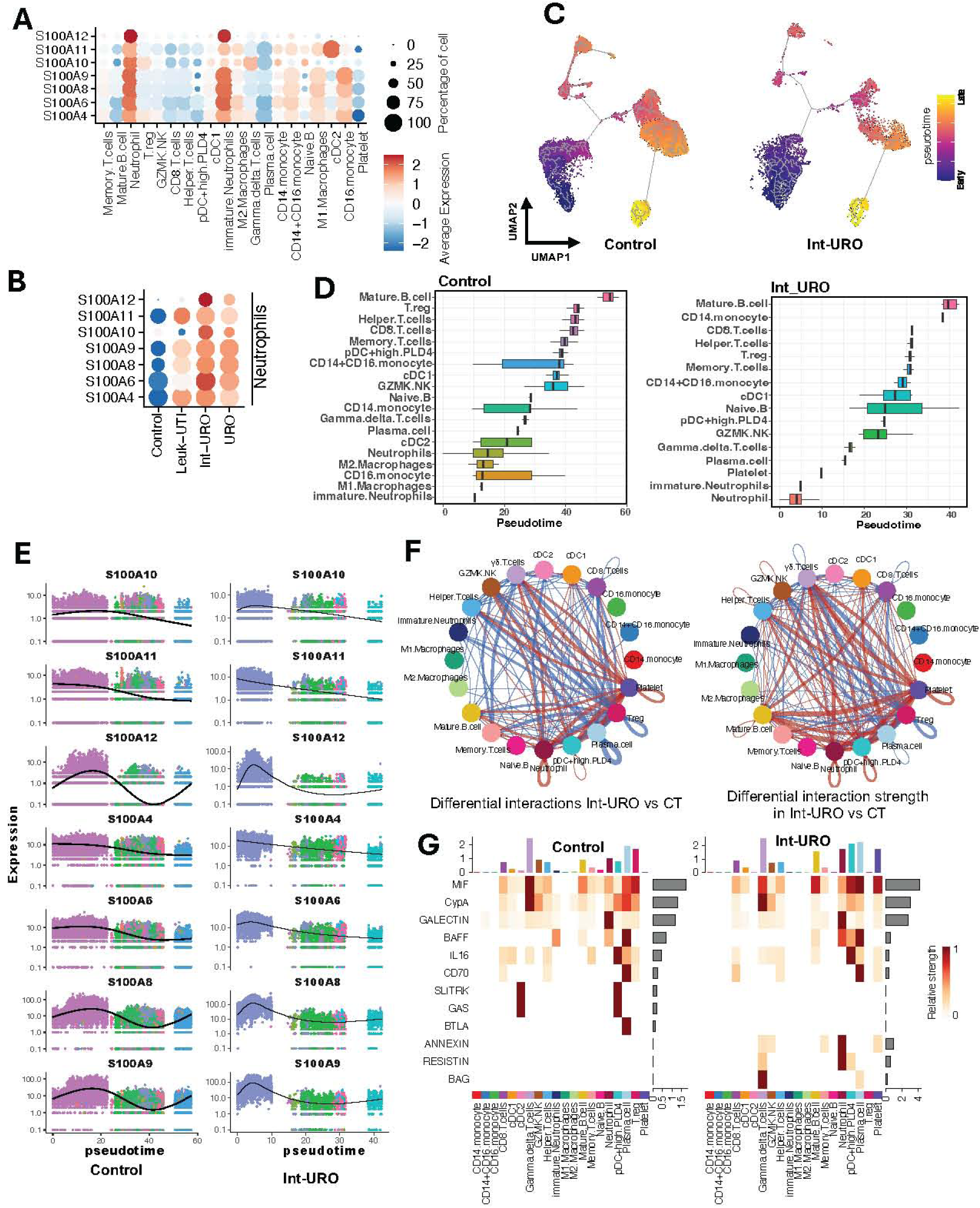
Pseudotime and cell-cell communications in adult MODS. (A) Expression of S100 genes identified in pediatric MODS across different cell types in adult MODS. S100 genes were highly expressed in neutrophils and immature neutrophils in adult MODS, consistent with the findings in pediatric MODS. (B) Expression levels of S100 genes in neutrophils across control, Leuk-UTI, Int-URO, and URO (post MODS) samples. S100 gene expression increased with disease progression, peaking at Int-URO. Blue indicates low expression, and red indicates high expression. The size of each circle represents the fraction of cells expressing S100 genes. (C) Pseudotime analysis of cells in control, and Int-URO. As the disease progresses, the abundance of immature neutrophils increases, reflected in the higher proportion of cells at early pseudotime stages. (D) Boxplot showing the median pseudotime of each cell type in control (left panel) and Int-URO (right). (E) Variations in S100 gene expression along pseudotime in control and Int-URO samples. S100 gene expression exhibits diverse patterns with pseudotime progression in both groups. (F) Differential cell-cell communication networks in Int-URO compared to control. The lower panel shows interaction strength, with thicker lines indicating stronger interactions, calculated based on significant co-expression of ligand-receptor pairs (log_2_ fold change ≥ 0.5, p-value ≤ 0.05). (G) Overall signaling strengths in individual cell types in control and Int-URO.

We further conducted pseudotime analysis to assess the differentiation status of cells. In controls, neutrophils displayed advanced maturation along with other monocytes and macrophages (Fig. 7C-D). In contrast, neutrophils in Int-URO exhibited greater stemness, as indicated by dense blue clusters in the pseudotime (Fig. 7C). Furthermore, Int-URO demonstrated early maturation of plasma cells, gamma delta T cells, GZMK.NK cells, and naïve B cells, a pattern not observed in control samples (Fig. 7D).

Analysis of *S100* gene expression along pseudotime revealed high initial expression in neutrophils, which decreased over time in controls and Int-URO, except for *S100A8, S100A9,* and *S100A12* (Fig. 7E). These genes demonstrated an oscillatory expression pattern, with the highest oscillation observed in controls, followed by Int-URO. These findings suggest that while neutrophil abundance and *S100* gene regulation during MODS initiation are similar in adult and pediatric MODS, the broader cellular landscape in adult MODS is adversely affected, with a lack of significant inflammatory responses.

Cell-cell communication analysis revealed enhanced interactions in adult MODS (Int-URO), particularly among neutrophils, pDC+high PLD4 cells, mature B cells, Vδ T cells, CD8+ T cells, and helper T cells (Fig. 7F), and major source of signaling in Int-URO were mature B cells, neutrophils and pDC+highPLD4 cells (Figure S11A). Notably, BAG, RESISTIN, and ANNEXIN signaling pathways were specifically activated in neutrophils and Vδ T cells in adult MODS, but absent in pediatric MODS (Fig. 7G and Figure S11B). These findings highlight differences in signaling between adult and pediatric MODS, despite a few shared characteristics that may provide insights into the core molecular mechanisms underlying MODS.

## Discussion

Despite decades of research, drug development for sepsis using both animal and translational models has largely failed to yield significant clinical benefits. In our previous study we demonstrated that neutrophils are the primary cells driving gene expression in pediatric MODS^12^. This led us to conduct the first comprehensive single cell RNA-seq analysis of pediatric patients developing MODS, which shows the complexity and heterogeneity of gene expression across all circulatory immune cells. We also compared our pediatric data with adult MODS (ICU patients at the onset of MODS) to gain deeper insights into the molecular mechanisms underlying this condition.

Our study revealed that circulatory immune cells in pediatric MODS patients exist in a state of “functional anarchy” characterized by the simultaneous upregulation of hyperinflammatory gene expression patterns and the downregulation of immune responses. Comparative analysis of ICU patients at the onset of MODS in pediatric and adult populations highlighted both key differences and similarities in immune cell behavior. Enhanced neutrophil activity and reduced pDC+highPLD4 cells were observed in both groups of MODS patients. However, adult MODS exhibited broader immune cell dysregulation compared to pediatric samples, which may be attributed to the more mature immune system in adults compared to children ^27,28^. In both adult and pediatric MODS, immune cells demonstrated decreased protein synthesis and increased apoptotic activity, likely as mechanisms to maintain immune homeostasis and reduce metabolic demand. Notably, only pediatric MODS showed enhanced inflammasome activity across various cell types.

The inflammasomes is a multiprotein complex that is responsible for the innate immune response to microbes that regulates the activation of caspases and inducing inflammation resulting in pyroptosis ^29^. Increased inflammasome activity was observed in pDC+high PLD4 cells, mature B cells, CD8+ T cells, GZMK+ NK cells, helper T cells, and memory T cells underscores the critical role of the inflammatory response during the early stages of MODS in pediatric patients. This pro-inflammatory state was further evidenced by the upregulation of crucial genes, including *IL32, MIF, RASA3*, and *CCR3,* all of which are strongly associated with inflammation. Notably, *IL32* is a well-characterized pro-inflammatory cytokine that induces the production of other cytokines, such as tumor necrosis factor (TNF)-α, *IL-6*, and *IL-1β* ^30^. The enhanced inflammatory response in pediatric MODS included upregulation of several pro-inflammatory regulons, including IRF5(+), XBP1(+), YBX1(+), MYB(+), and PAX5(+) ^31,32^. In contrast, neutrophils and Treg cells simultaneously exhibited immune cell activation including *CD27, CD38,* and *CD69,* while showing downregulation of several inflammasome-related genes, such as *JUNB, NFKBIA, FOSB, FOS, IL1B,* and *HLA.* These observations highlight neutrophils as central regulators of inflammation, modulating critical genes, including members of the *S100* gene family.

The *S100* family of genes, which mediate multiple protein pathways, are situated adjacently on the largest chromosome 1 in humans. These genes play a role in maintaining immune and cellular homeostasis by acting through multiple pathways. The *S100* genes are a broad subfamily of low-molecular weight calcium-binding proteins with structural similarity and functional discrepancy ^33^. It’s just as important for the immune system to be able to tolerate beneficial bacteria such as intestinal microbiota or probiotics as recognize harmful ones. In pediatric MODS, we identified *S100A4, S100A6, S100A8, S100A9, S100A10, S100A11,* and *S100A12* which function as DAMPs ^17,18,34^, were specifically expressed in neutrophils, and these genes has been associated with increased neutrophil infiltration into tissues and organs ^35,36^. In one adult MODS cohort, the expression levels of *S100* genes were significantly increased during the first 72 hours of hospitalization in MODS patients, a pattern absent in controls and non-MODS patients (Figure S4C), indicating MODS-specific regulation. The consistent early upregulation of these *S100* genes in adult MODS further supports their importance in MODS pathophysiology. These genes primarily promote inflammation in pediatric patients, they might be playing other roles in adults. Pseudotime analysis revealed distinct dynamics of *S100* gene expression between adult and pediatric MODS, suggesting the multifunctional roles of *S100* genes in both groups. This variability may explain the distinct contributions of *S100* genes to early MODS in pediatric patients, where they play a critical role, compared to their other roles in adult MODS, highlighting their context-dependent involvement in MODS pathogenesis. Despite belonging to the same gene family, each *S100* gene exhibits distinct expression and distribution patterns across tissues and cell types. Among these, *S100A4, S100A8, S100A9, S100A10, S100A12, S100A14, S100P,* and *S100Z* are specifically expressed in immune cells ^37^. In our study, the identified *S100* gene cluster in pediatric MODS demonstrated distinct performance across multiple cohorts, highlighting the potential as diagnostic biomarkers for the loss of inflammatory homeostasis. Furthermore, these *S100* genes represent promising therapeutic targets for pediatric MODS. Drugs such as Paquinimod, Laquinimod, Tasquinimod, Cromolyn, Niclosamide, and SEN205A, along with neutralizing antibodies targeting specific S100 proteins (S100A4, S100P, S100A9) ^38–42^, could potentially be repurposed to restore immune homeostasis and prevent MODS ^43^.

Pathways designed to dampen the adaptive immune response were also found to be activated. Mechanisms such as restricted protein synthesis and reduced expression of surface proteins, including HLA-DRB5, HLA-DQA2, and IgG, which are critical for antigen processing were affected ^44,45^. These processes could potentially contribute to a dysregulated immune environment in response to antigens and pathogens. Further evidence for this dysregulation was provided by cell-cell communication analysis, which identified TNFSF13, and Gal-9 (Lgals9) ligand-receptor pairs involved in cell-cell interactions. These molecules are associated with immune suppression and T cell apoptosis ^46,47^. While pediatric MODS shares some of the same response characteristics with adult MODS, adults exhibit a more robust immune response during the early MODS. This underscores the need for large scale multicenter studies to explore these pathways and improve understanding of how sepsis transitions into MODS in both adult and pediatric patients.

Currently, two nested NIH-sponsored studies are investigating a dual layered therapeutic approach for sepsis-induced MODS. The GRACE-2 trial focuses on treating immune-paralysis in pediatric MODS with granulocyte macrophage-colony stimulating factor (GM-CSF), while the TRIPS trial evaluates the use of anakinra (or placebo) to reverse systemic inflammation in pediatric MODS ^48^. However, our findings suggest that at the onset of pediatric MODS, both hyper-inflammatory and immune-suppressive phenotypes coexist in the same patient indicating that neither approach by itself may be adequate to effectively treat these patients.

The limitation of our study is the small number of pediatric patients in each group and the variability in their ages. It is important to note that these pediatric MODS patients are uncommon, and capturing the timing of transition sometimes proved challenging and excluded some potential subjects. It took us more than two years to capture this cohort in one of the largest children’s hospitals in Michigan, highlighting the significance and difficulty involved in conducting this study. This underscores the need for future studies to generate data from multiple centers to develop deeper insights into the initiation and progression of MODS in pediatric patients and explore why some ICU patients develop MODS while others do not.

In conclusion, this study is the first to characterize immune cells using scRNA-seq in pediatric MODS patients at the point of clinical recognition and to compare their immune response with that of adult MODS patients. Pediatric MODS patients showed some overlap with adults, who, perhaps due to a lifetime of antigen exposure, exhibit a stronger immune response. The data and insights provided here offer valuable resources for understanding the progression from ICU sepsis to MODS. Our findings suggest that insights from adult MODS can be leveraged to enhance the understanding of MODS initiation and progression in pediatric patients. Upstream influences such as *S100* genes which may have a role in maintaining immune homeostasis, could serve as diagnostic markers and therapeutic targets.

## Material and Methods

### Ethics

The Institutional Review Board (IRB) approval for this study (2019-113-SH/HDVCH) was obtained from Corewell Health on January 13, 2019. As all participants were minors and incapacitated by critical illness, consent was obtained from their parents prior to recruitment. Only de-identified patient samples and data from the PICU at Helen DeVos Children’s Hospital were used in this research.

### Patients and blood samples

Pediatric patients who presented with sepsis in the ICU were followed carefully and on meeting MODS criteria were recruited within 24 hours into the study. Parameters used in the Age of Blood in Children in Pediatric Intensive Care Unit (ABC-PICU) trial ^49^. PODIUM criteria were used afterwards to confirm systems affected ^50^. After confirmation, patients were recruited, and blood were obtained. Blood samples were collected only once within 24 hours of identifying the patients with MODS (N=5). Blood samples were drawn into EDTA tubes, stored, and then transported immediately at 4-8°C to a processing laboratory. Control patients (N=5) who received sedation for brief same day procedures were included. Nucleated cells from blood samples were separated within 4 hours. The control individuals were age and biological sex-matched with the MODS patients.

### Flow cytometry analysis to capture immune cells

Cells isolated from the EDTA blood collection using a AutoMACS cell separator (Miltenyi) were washed in PBS and pelleted at 300 g for 5 minutes. Cell pellets were resuspended in staining buffer (BD Biosciences, San Jose, CA) and manually counted. After counting, cells were subjected to Fc blocking (Biolegend, cat#422302) for 10 min. After Fc blocking, cells were labeled with two separate panels as follows: CD45-APC, CD3-APC-R700, CD19-FITC, CD4-BV711, CD8-Pacific Blue, CD25-PE CD69-APC-Cy7, CD45ra-BV510, CD45ro-BV605, PD1-BV650, and IgD-BV421; or CD45-APC, CD11b-Pacific Blue, CD11c-BV650, HLA-DR-APC-Cy7, CD80-BV711, CD64-FITC, PD-L1-BV421, CD56-APC-R700, CD206-BV510, CD14-BV605, and CD16-PE. After 30 min incubation at 4°C, cells were washed and resuspended in 200 µL of buffer containing 7-AAD (BD Biosciences, cat#559925). Fluorescence was detected and quantified using a ZE5 flow cytometer (BioRad, Hercules, CA), and analyzed by FlowJO software (BD Biosciences).

### Single cell RNA sequencing

Sorted CD45-positive cells were processed for scRNA-seq using the 10X Chromium 5’ Library and Gel Bead Kit from 10x Genomics. Following GelBead in-Emulsion reverse transcription (GEM-RT), 12–15 cycles of PCR amplification were conducted to obtain cDNAs for RNA-seq library preparation. Libraries for individual samples were prepared according to the manufacturer’s instructions and sequenced on an Illumina NovaSeq 6000 Sequencing System. Library preparation and sequencing were performed in the Genomics Core at Van Andel Institute.

### Adult MODS scRNA-seq data

We obtained publicly available adult scRNA-seq data and corresponding sample metadata ^26^ from the Broad Institute’s Single Cell Portal. This dataset includes scRNA-seq data from 65 patients, categorized as follows: controls (n=19), urinary-tract infection (UTI), with leukocytosis (blood WBC ≥ 12,000 per mm3) but no organ dysfunction (Leuk-UTI) (n=10), UTI with mild or transient organ dysfunction (Int-URO) (n=7), UTI with clear or persistent organ dysfunction (URO) (n=10), bacteremic individuals with sepsis in hospital wards (Bac-SEP) (n=4), intensive care unit (ICU) patients with sepsis (ICU-SEP) (n=8), and ICU patients without sepsis (ICU-NoSEP) (n=7). The scRNA-seq data were generated from the CD45+ cell population, and Int-URO, the group similar to pediatric MODS, was specifically studied.

### Processing and analysis of scRNA-seq data

The FASTQ sequencing reads were mapped to the Hg38 transcriptome using the ENSEMBL GRCh38.p3 annotation with Cell Ranger count (version 6.1.1). The --force-cells=10k and --include-introns options were employed to generate the filtered matrix files as suggested by 10x Genomics for studies enriched with neutrophils ^15^. The data can be accessed at Gene Expression Omnibus (GEO) under accession number GSE269751. The filtered matrix files were imported into the Seurat R package (version 4.1.2) for downstream analysis. During the quality control, cells with gene counts between 200 and 2500 and mitochondrial gene percentage below 25% were retained. Expression data was log normalized, followed by the selection of the top 5,000 variable genes for scaling. All the samples were integrated together to create a single Seurat object. Further, a commonly used Seurat workflow was adopted to process the data. Clustering of the cells was performed with 0.8 resolution. Cells were manually annotated based on their expression of canonical marker genes provided by the Human Cell Atlas.

### InferCNV for gene cluster identification

Cells with cluster of enhanced gene expression were identified using InferCNV ^16^ (https://github.com/broadinstitute/inferCNV) with default parameters. InferCNV originally developed to infer CNVs from gene expression in tumor cells, could reveal distinct clusters shared by individual MODS patients. InferCNV estimates CNVs by analyzing consistent over- or under-expression of genes in a genomic region relative to a reference, uses a sliding window approach across the genome to smooth expression data, and highlights regions where significant deviations from the reference occur, which can further help to identify cluster of genes overexpressed in each cell. All the MODS cells were used as input, and cells from the control group were used as a reference. We identified the cells with the extent of cluster of over expressed genes in each cell type in individual MODS patients.

### Cell deconvolution in bulk RNA-seq data

Normalized gene expression profiles and cell type markers were utilized to compute single-sample gene set enrichment analysis (ssGSEA) scores using the sigatlas package (https://github.com/Bin-Chen-Lab/sigatlas). An ssGSEA score represents the enrichment of each cell type in a target sample, with a higher enrichment score indicating a greater abundance of that cell type. To minimize technical variation among samples, we adopted the same pipeline to process the raw data and made comparisons within the same cohort. Geometric mean of the expression of a gene set was calculated for each sample and further used to re-classify patients from disease groups. Receiver operating characteristic (ROC) and area under curve (AUC) implemented in the pROC package ^51^ were used to assess the performance. Multiple independent datasets were used as validation cohorts. For those published datasets, we used normalized gene expression counts from publicly available RNA-seq data and adopted the pipeline from our previous study ^52^ to process the GEO microarray data.

### Identify cells with active genes in scRNA-seq data

For gene set enrichment analysis related to inflammasome, apoptosis, and immune cell activation, we obtained the gene lists from the Molecular Signatures Database (MSigDB) (https://www.gsea-msigdb.org/gsea/msigdb/index.jsp). Subsequently, we used the AUCell package ^53^ to identify the cells with active gene sets. The enrichment threshold for each cell was defined as the greater value between 1 and (max_score - 1), where max_score represents the highest AUCell enrichment score achieved by any gene set within a specific cell. For cells enriched by multiple gene sets, the assignment was made to the gene set with the higher number of expressed genes (expression count > 0).

### Pathway Analysis

For each cell type, differential expression analysis between control and MODS was performed using the Wilcoxon rank sum test to identify differentially expressed (DE) genes. Genes with min PCT 25%, log_2_ fold change ≥0.5, and adjusted P-value ≤0.05 were considered as DE genes. ClusterProfiler ^54^ was used to perform the pathway analysis and ggplot2 was implemented to visualize enriched pathways.

### Pseudotime analysis

Cell trajectories were inferred using Monocle3^55^ with default parameters. We selected the cluster with highest expression of stemness genes as a starting point to order cells.

### Cell surface protein abundance prediction

We utilized SPIDER ^56^ to predict cell surface protein abundance from our scRNA-seq data. SPIDER is a zero-shot model trained on various CITE-seq (cellular indexing of transcriptomes and epitopes) datasets and can be applied to any scRNA-seq datasets for protein abundance prediction. Only high confidence predictions were selected for further analysis. We computed the differentially expressed proteins (DEPs) between MODS and control. The DEPs with log_2_ fold change ≥ 0.5 with adjusted p-value ≤ 0.05 were considered as upregulated, whereas DEPs with log_2_ fold change ≤ −0.5 with adjusted p-value ≤ 0.05 were considered as downregulated. We visualized the distribution of these genes using the ggplot2 package in R.

### Regulatory network inference

A single-cell regulatory network in scRNA-seq data was constructed with SCENIC ^53^. The python package pySCENIC (https://pyscenic.readthedocs.io/en/latest/#) was used to infer gene regulatory networks from raw count data. The potential direct-binding targets (regulons) were identified based on DNA-motif analysis and gene regulatory network activity in individual cells was estimated using this package.

### Cell-cell communication

The CellChat package ^57^ was employed to identify ligand-receptor pairs responsible for cell-cell communications. This package utilizes information on ligand-receptor interactions and related pathways sourced from CellChatDB (https://www.cellchat.org/). CellChatDB encompasses manually curated literature-based signaling molecule interactions, including complexes with multimeric ligands and receptors, as well as cofactors influencing these interactions such as soluble agonists, antagonists, co-stimulatory, and co-inhibitory membrane-bound receptors. The algorithm infers potential interactions of ligand-receptor pairs along with their intensity between two cell populations, based on gene expression in the cell population, signaling cofactors, and the percentage of cells expressing the ligand or receptor. The significance of an interaction is assessed through a permutation test. For our analysis, receptors and ligands expressed in more than 25% of cells in a specific cell type with a p-value ≤ 0.01 and log_2_ fold change ≥ 0.5 were included. The interactions were identified and quantified based on the differentially over-expressed ligands and receptors for each cell group (p-value ≤ 0.05). Outgoing and incoming signaling strengths were computed as the cumulative interaction intensities of ligands and receptors expressed in the respective cells.

## Data and Code Availability

The codes used in these analyses are available at https://github.com/Bin-Chen-Lab/MODS/tree/master/scRNA-seq. The scRNA-seq data generated in this study is available through NCBI GEO accession GSE269751.

## Acknowledgement

Research reported in this publication was supported by the National Institute Of General Medical Sciences of the National Institutes of Health under Award Number R01GM134307 and Eunice Kennedy Shriver National Institute of Child Health and Human Development of NIH under award number 1K99HD111575. The content is solely the responsibility of the authors and does not necessarily represent the official views of the National Institutes of Health. We would like to thank the Genomic Core facility at Van Andel Institute (VAI) for helping in scRNA-seq library preparation and sequencing.

## Author contributions

Conceived and designed the experiments: RS, BC, and SR. Performed the experiments: RS and AWG. Analyzed the data: RS, SP, SK, and RC. Wrote the paper: RS, BC, AWG, SR, and NH. Supervised the study: BC and SR. All authors read and approved the final version of the manuscript.

