## Supplementary information for "Comparative immuno-biology at clinical recognition of early multiple organ dysfunction syndrome in pediatric and adult patients using single-cell transcriptomics"

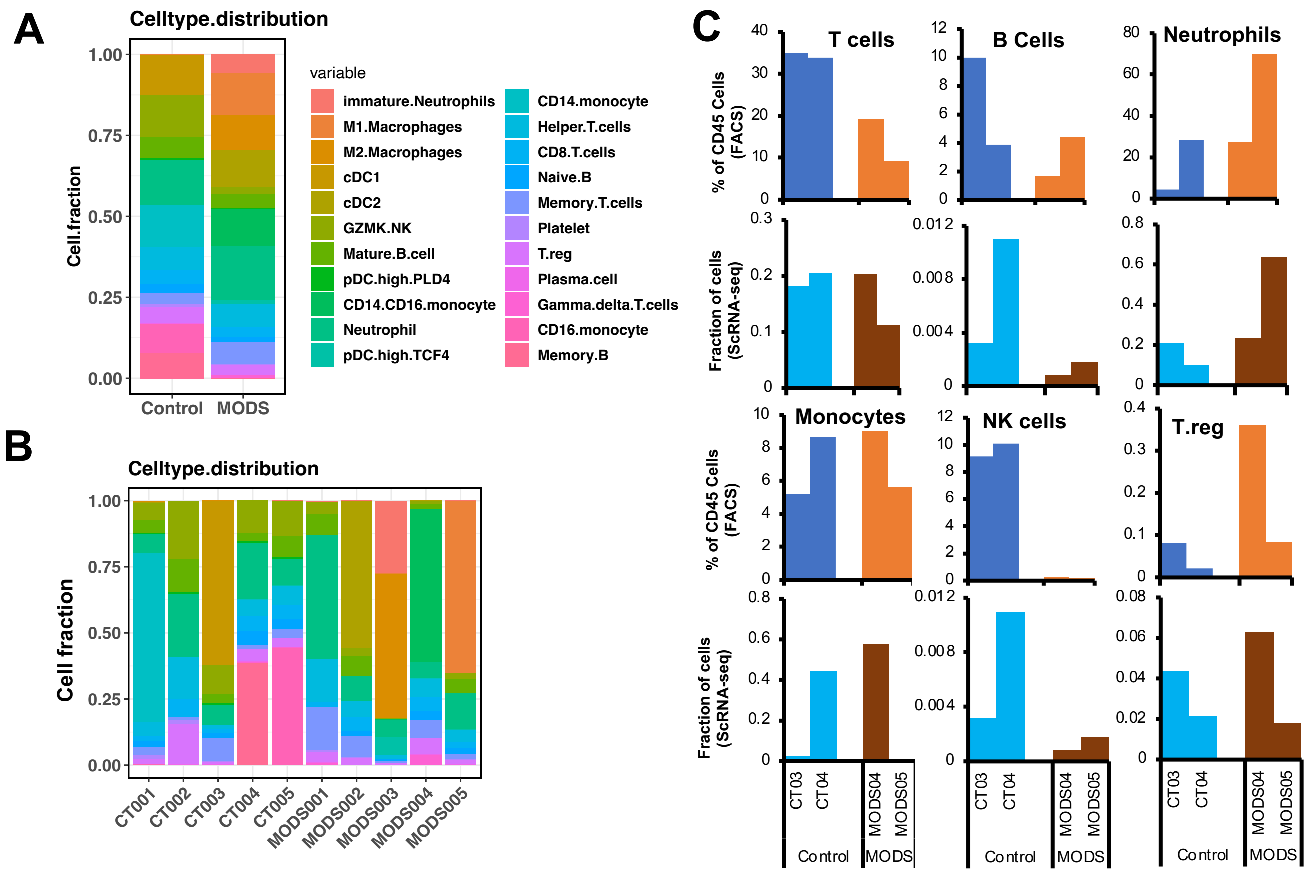

**Figure S1 Cell fraction distribution in control and MODS.** (A-B) Cell fraction distribution in control and MODS (A) and individual samples (B). (C) Comparison of cell fraction obtained from scRNA-seq and flow cytometry analysis.

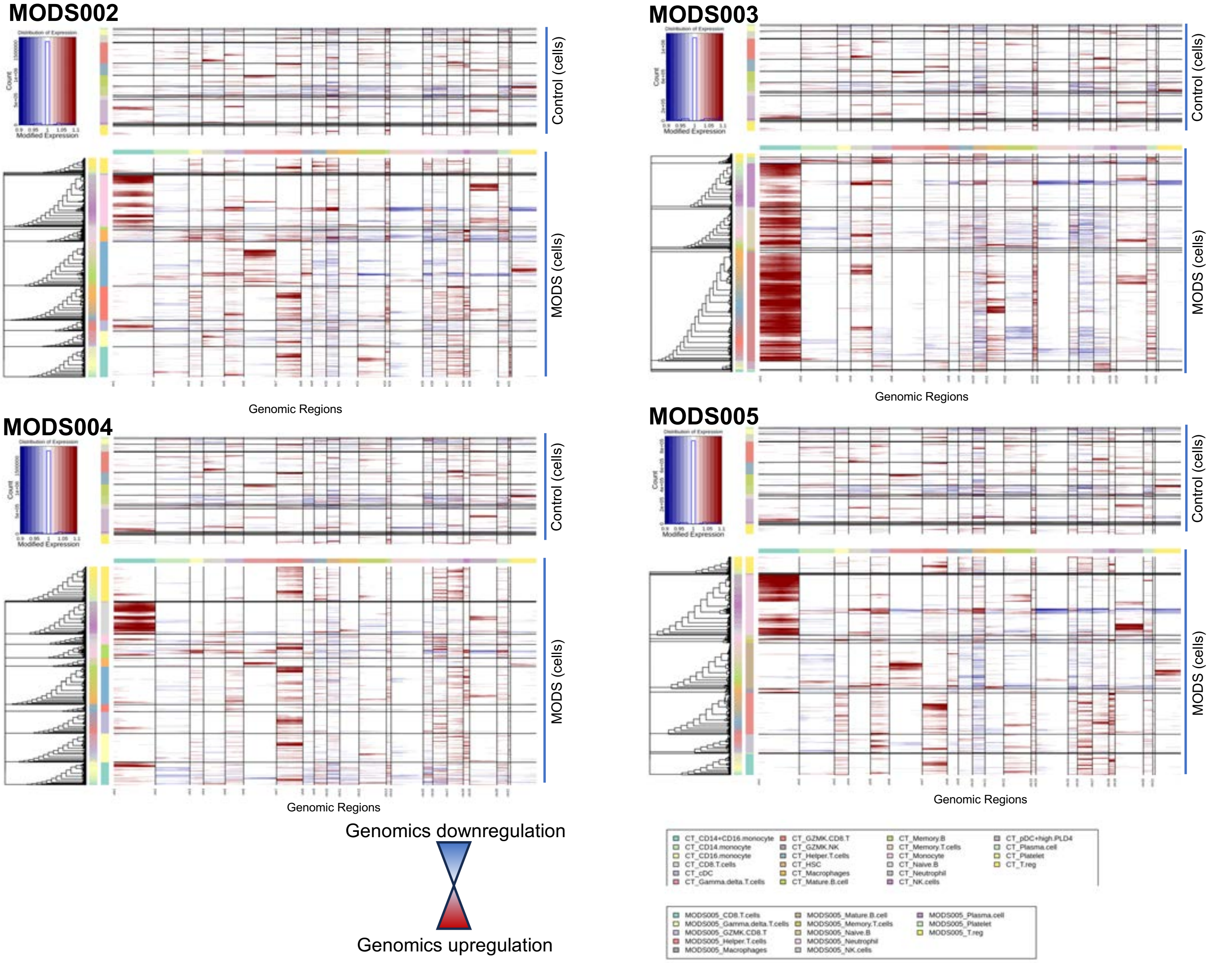

**Figure S2: InferCNV plots of all the cell types in control (upper) and MODS in individual patients.** The red color shows the amplified region, whereas blue lines shows the amplification loss in MODS compared to control. The the amplification of regions were specific to chr1 specifically in neutrophils.

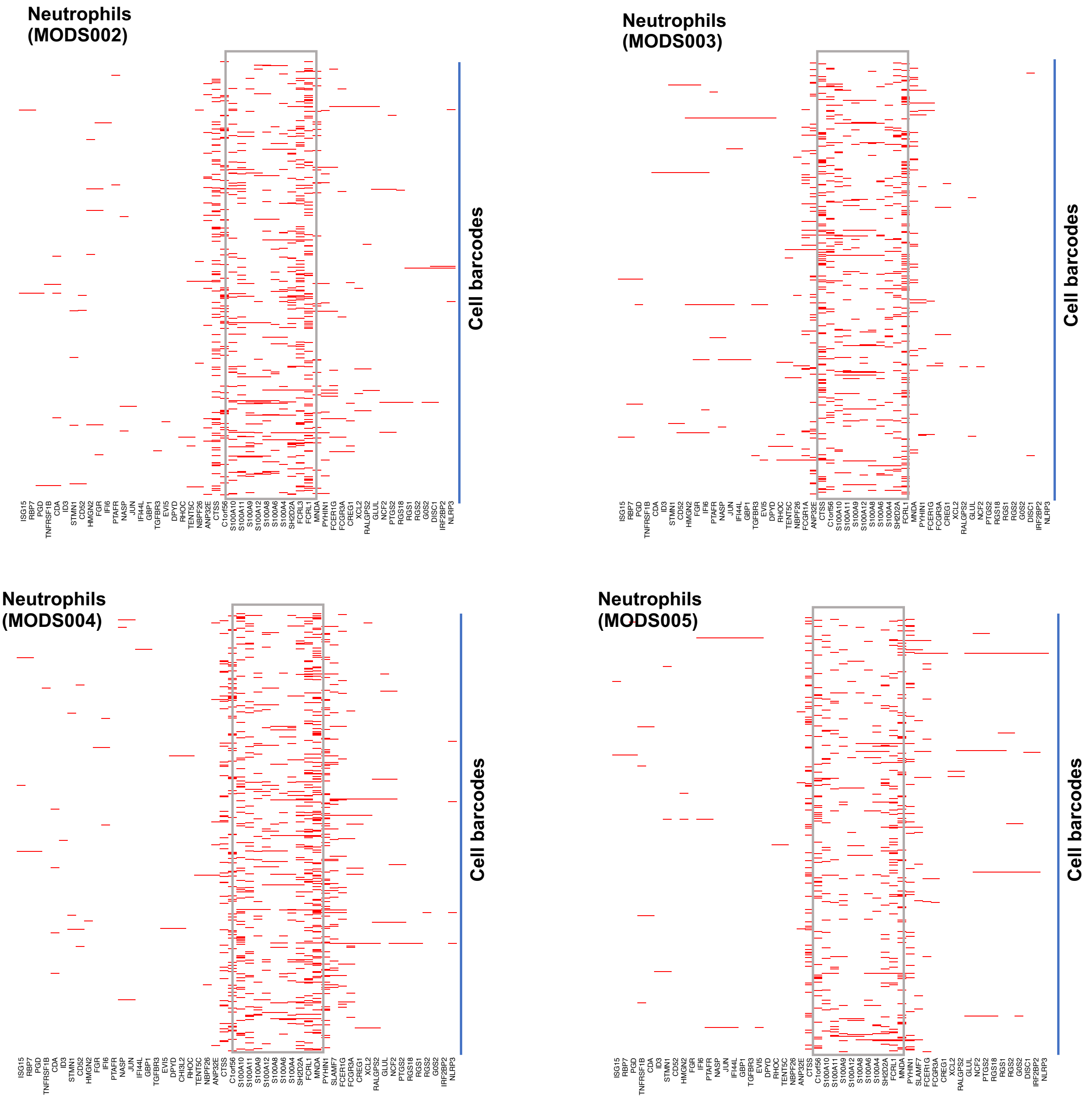

**Figure S3: Amplification of genes in neutrophils in individual MODS patients. The red color shows the amplification of genes in each cell.**

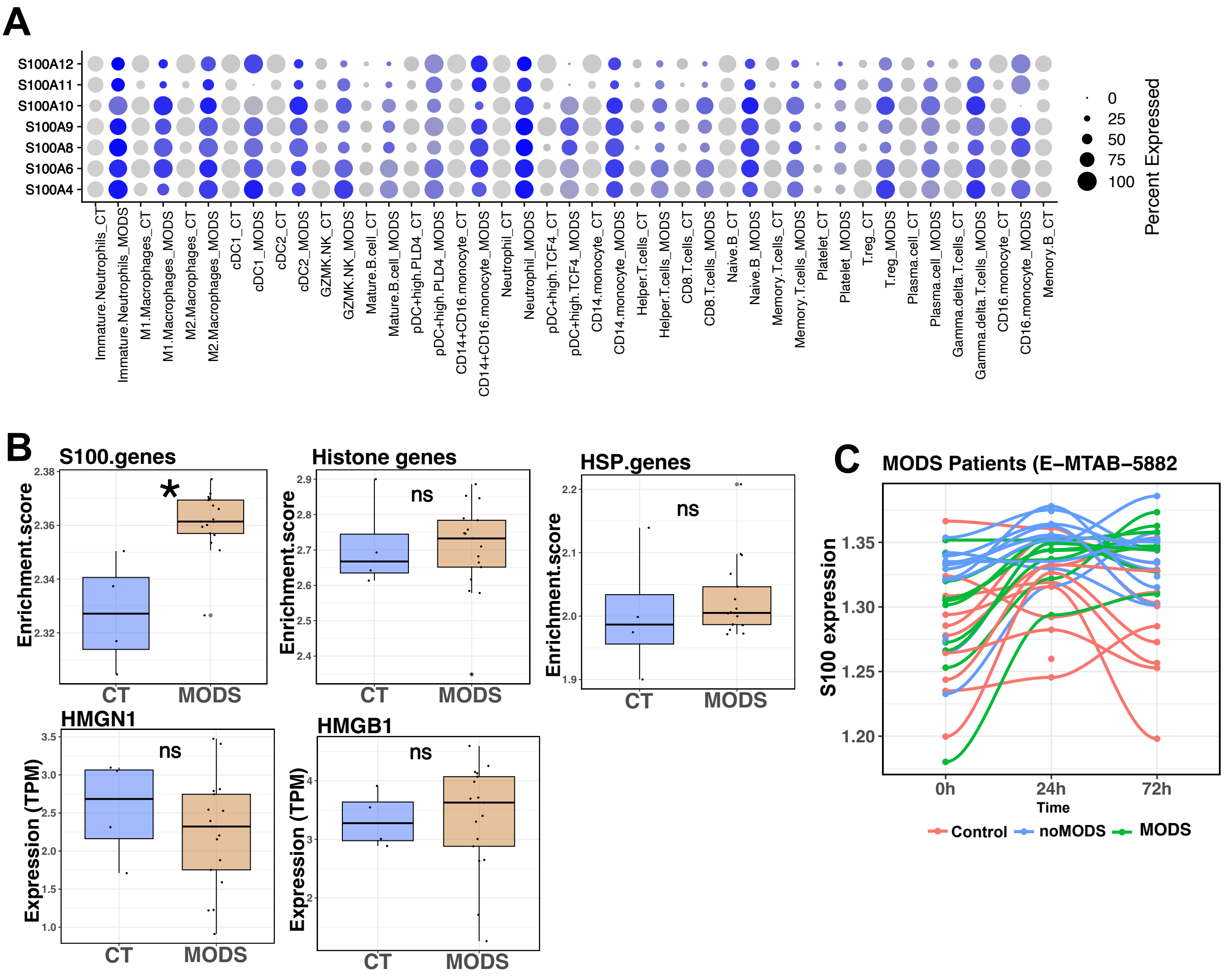

**Figure S4: Expression of S100 genes in control and MODS.** (A) Expression of S100 genes in different cell types in scRNA-seq. (B) Expression of other DAMP genes (Histone, HSP, HMGB1, and HMGN1) in control and MODS samples in the bulk RNA-seq data. None of these genes exhibited any change in expression between control and MODS. (C) Variation of S100 gene expression throughout different time points in control, and trauma patients without MODS (noMODS) and with MODS. **** 1e-16< p-value ≤ 1e-05, *** 1e-5 < p-value ≤ 0.0001, ns- not significant.

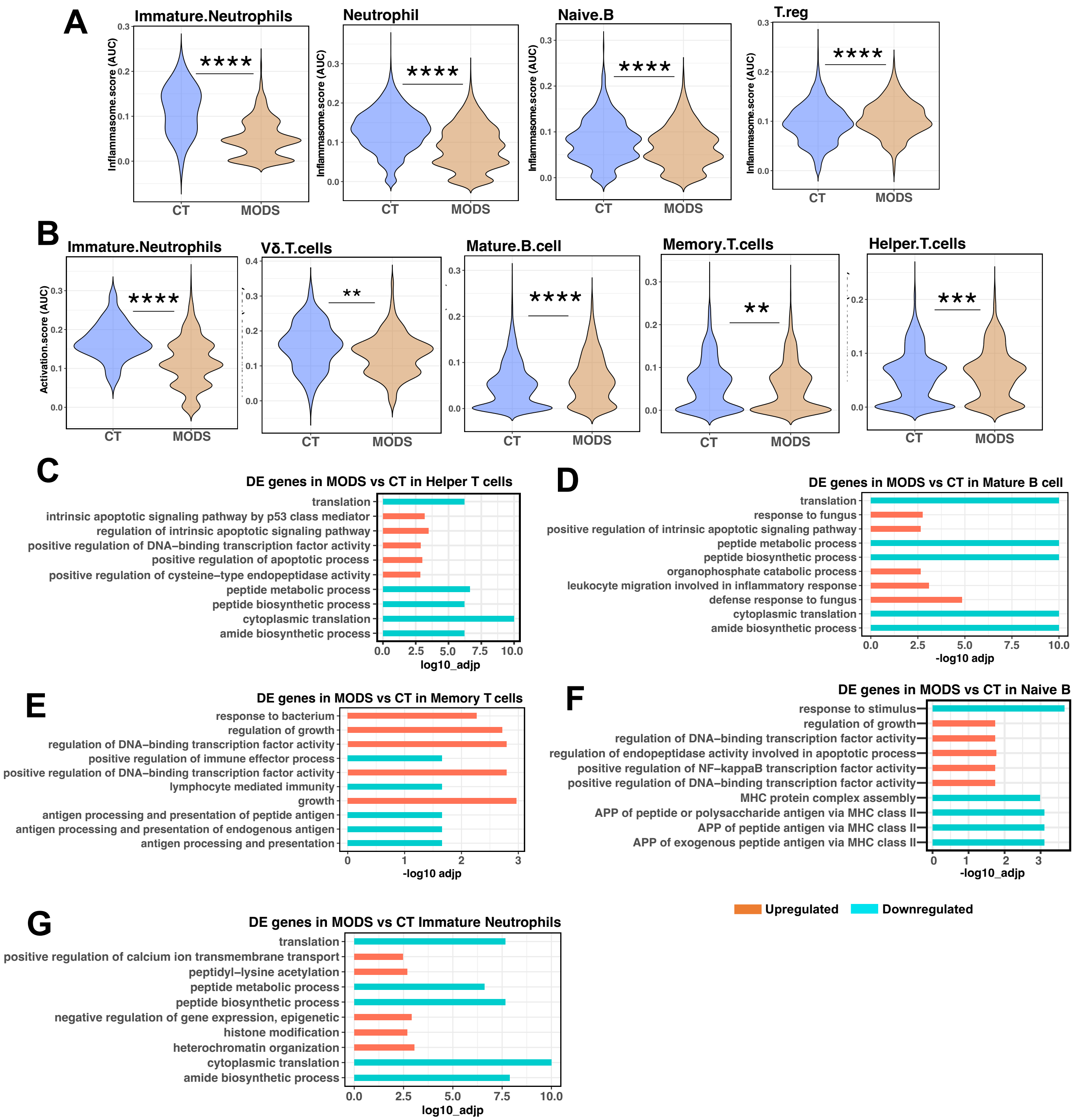

**Figure S5 Immune cells activity and enriched pathways in MODS.** (A) Inflammasome activity in immature neutrophils, neutrophils, naïve B cells, and Treg cells. (B) Immune activation activity in immature neutrophils, Vδ T cells, mature B cells, memory T cells, and helper T cells. All these cell types exhibited decreased immune activity in MODS. (C-G) Enriched biological processes identified by upregulated (red) and downregulated (blue) genes by comparing helper T cells (C), mature B cells (D), memory T cells (E), naïve B cells (F), and neutrophils (G) from MODS with control. Significance levels: **** 1e-16< p-value ≤ 1e-05, *** 1e-5 < p-value ≤ 0.0001, and ** 0.0001 < p-value ≤ 0.01.

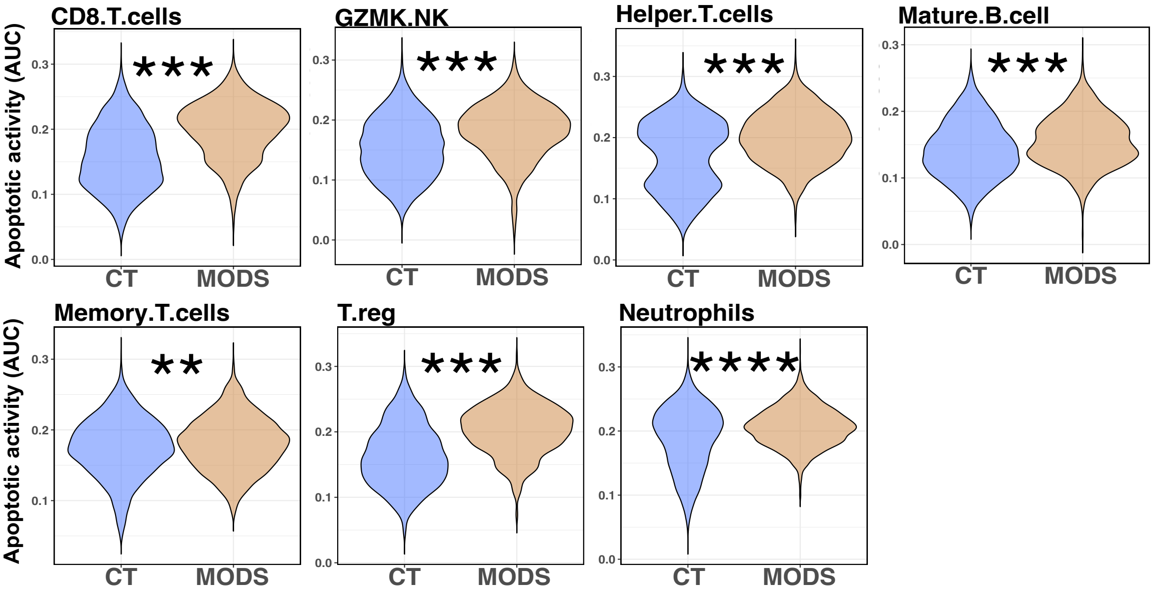

**Figure S6 Apoptotic activity of immune cells in control and MODS.** Most of the immune cells associated with adaptive immune response and neutrophils were showing the greater apoptotic activity in MODS. **** 1e-16< p-value ≤ 1e-05, *** 1e-5 < p-value ≤ 0.0001, and ** 0.0001 < p-value ≤ 0.01.

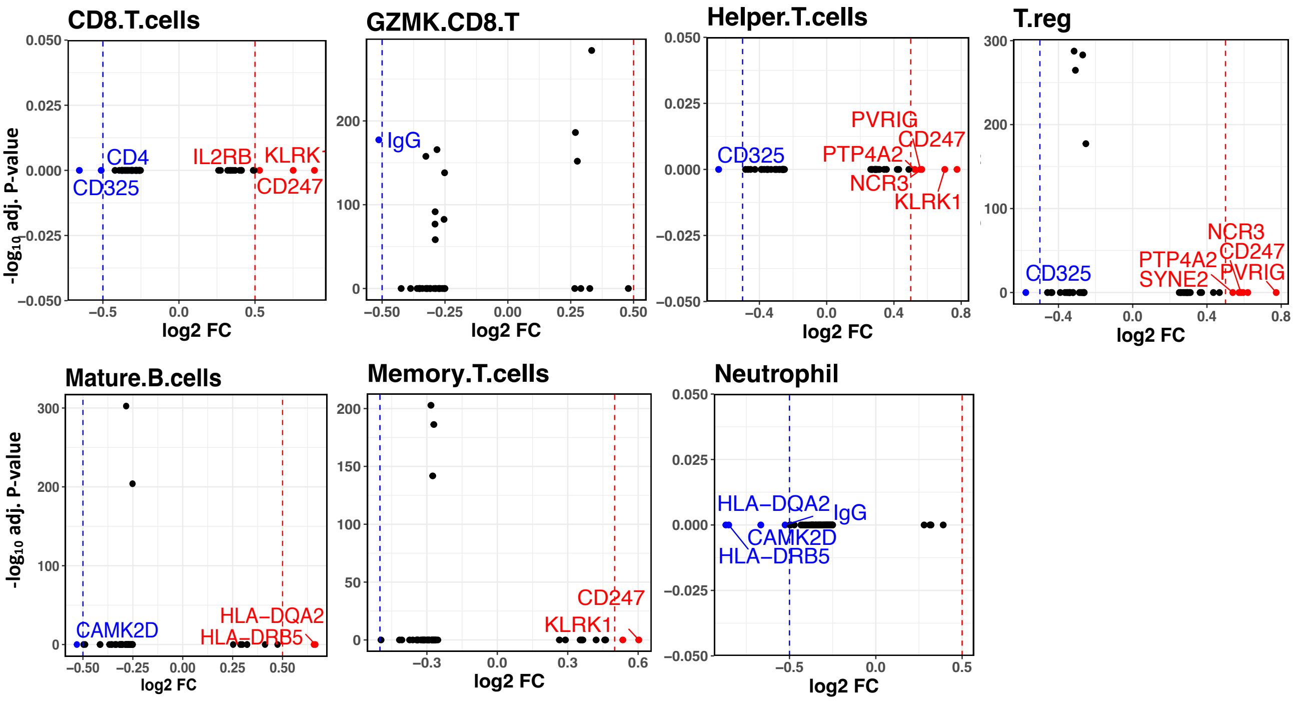

**Figure S7 Surface protein expression in immune cells.** Surface protein expression was analyzed using the SPIDER package (https://github.com/Bin-Chen-Lab/spider). Only high-confidence predictions with an absolute log2 fold change ≥ 0.5 and an adjusted p-value ≤ 0.05 were considered. CD8+ T cells, helper T cells, Treg cells, mature B cells, and memory T cells exhibited both upregulated and downregulated surface protein expression, whereas GZMK+ NK cells and neutrophils primarily showed downregulation of surface proteins.

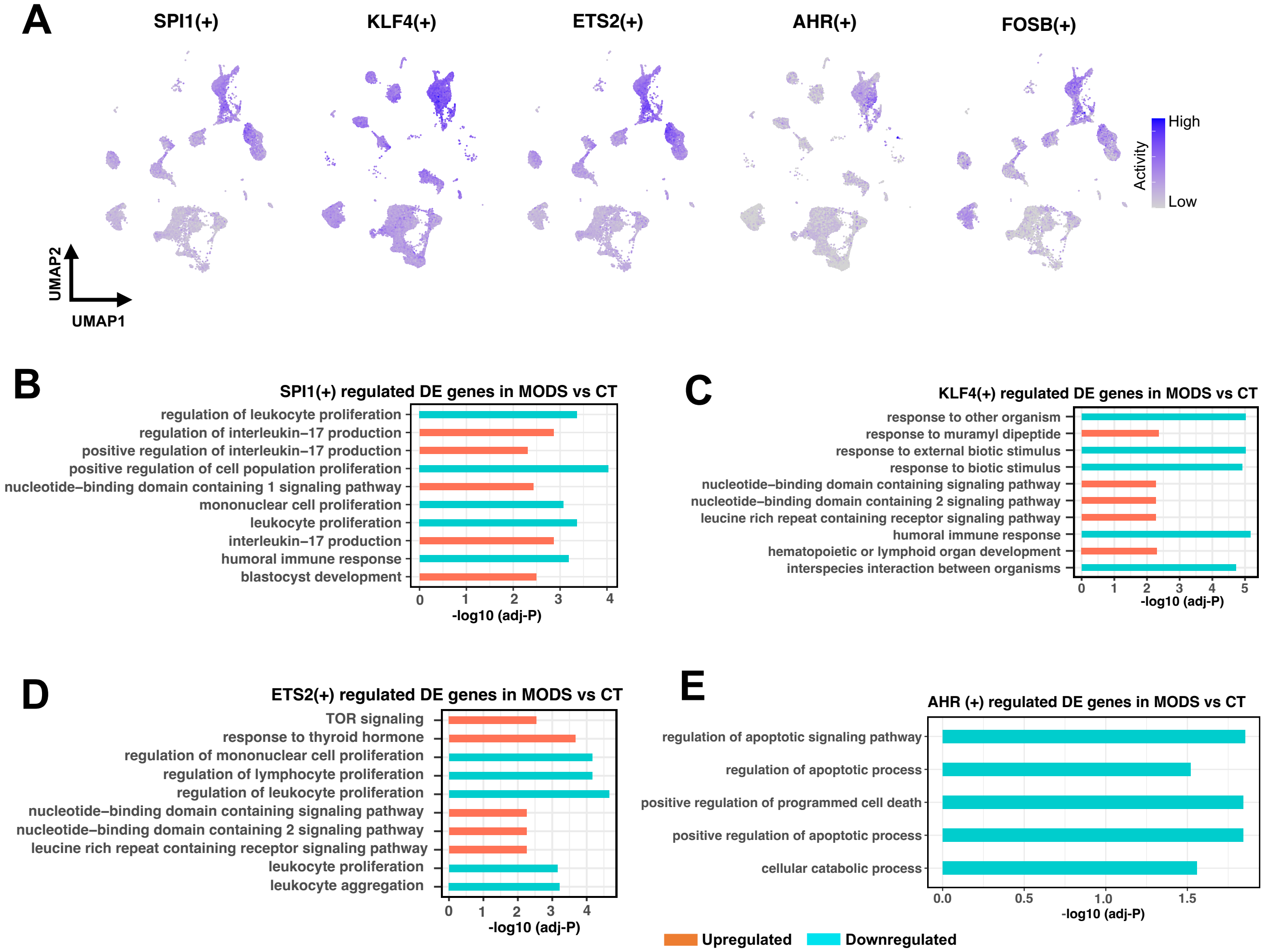

**Figure S8 TF activities in different cell types.** (A) Feature plot showing differential transcription factor (TF) activities in control and MODS. (B-E) Enriched biological processes associated with genes regulated by SPI1+ (B), KLF4+ (C), ETS2+ (D), and AHR+ (E) regulons in MODS (red) and control (blue).

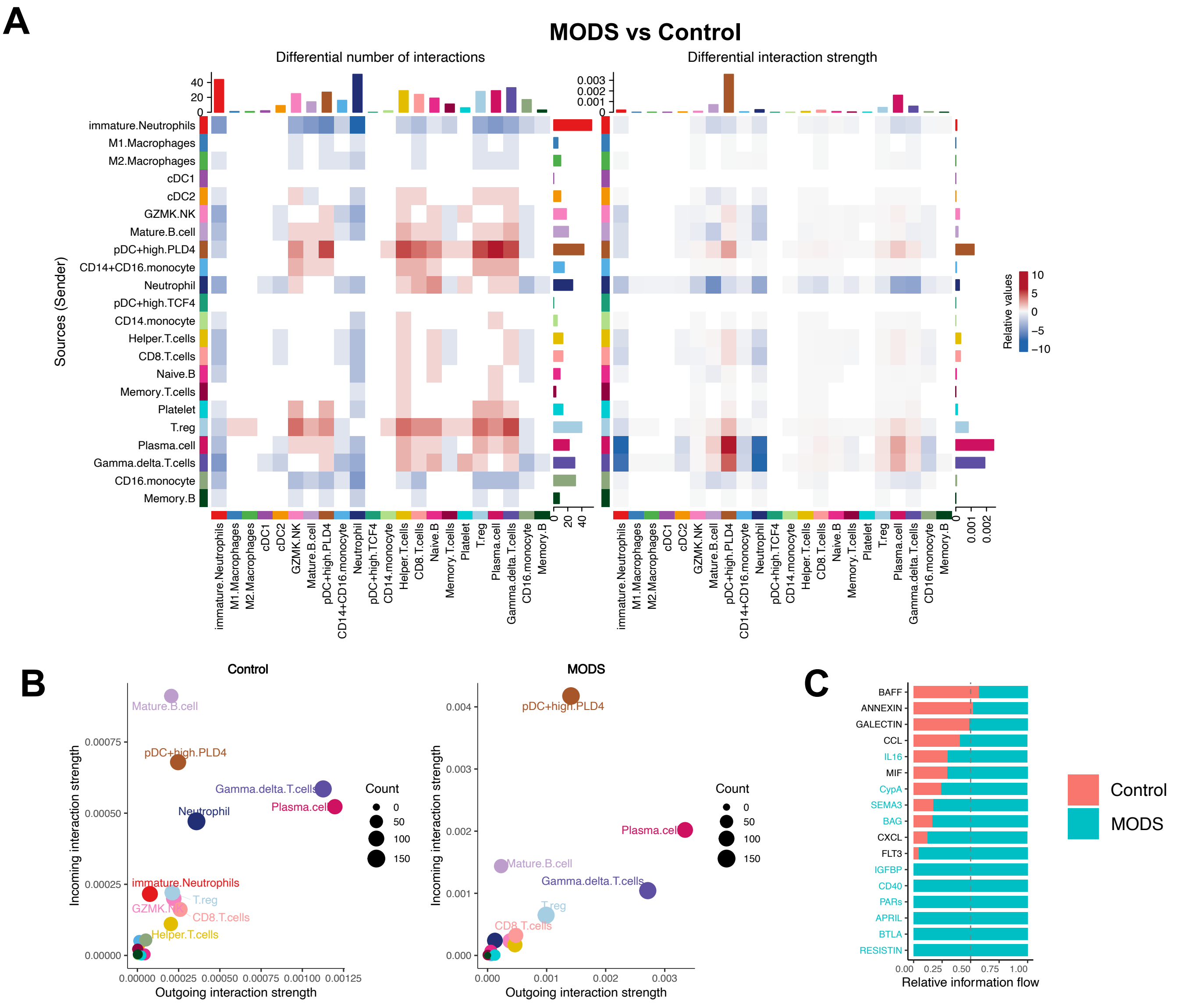

**Figure S9: Cell-cell communications among all cell types.** (A) Heatmaps displaying the differential interactions (left) and strength of interactions (right) in MODS compared to control. The red color indicates enhanced probability of interactions, while blue color represents decreased cell interactions. Enhanced interaction strength is observed in pDC+highPLD4 cells, plasma cells, and Vδ T cells in MODS compared to control. (B) Dot plot illustrating the key cell types involved in the majority of interactions in control and MODS. In control, B cells are the major cell type, whereas in MODS, pDC+highPLD4 cells are the major cell type associated with cell-cell communications. (C) Bar plot showing the significant signaling molecules involved in cell-cell communications in control and MODS.

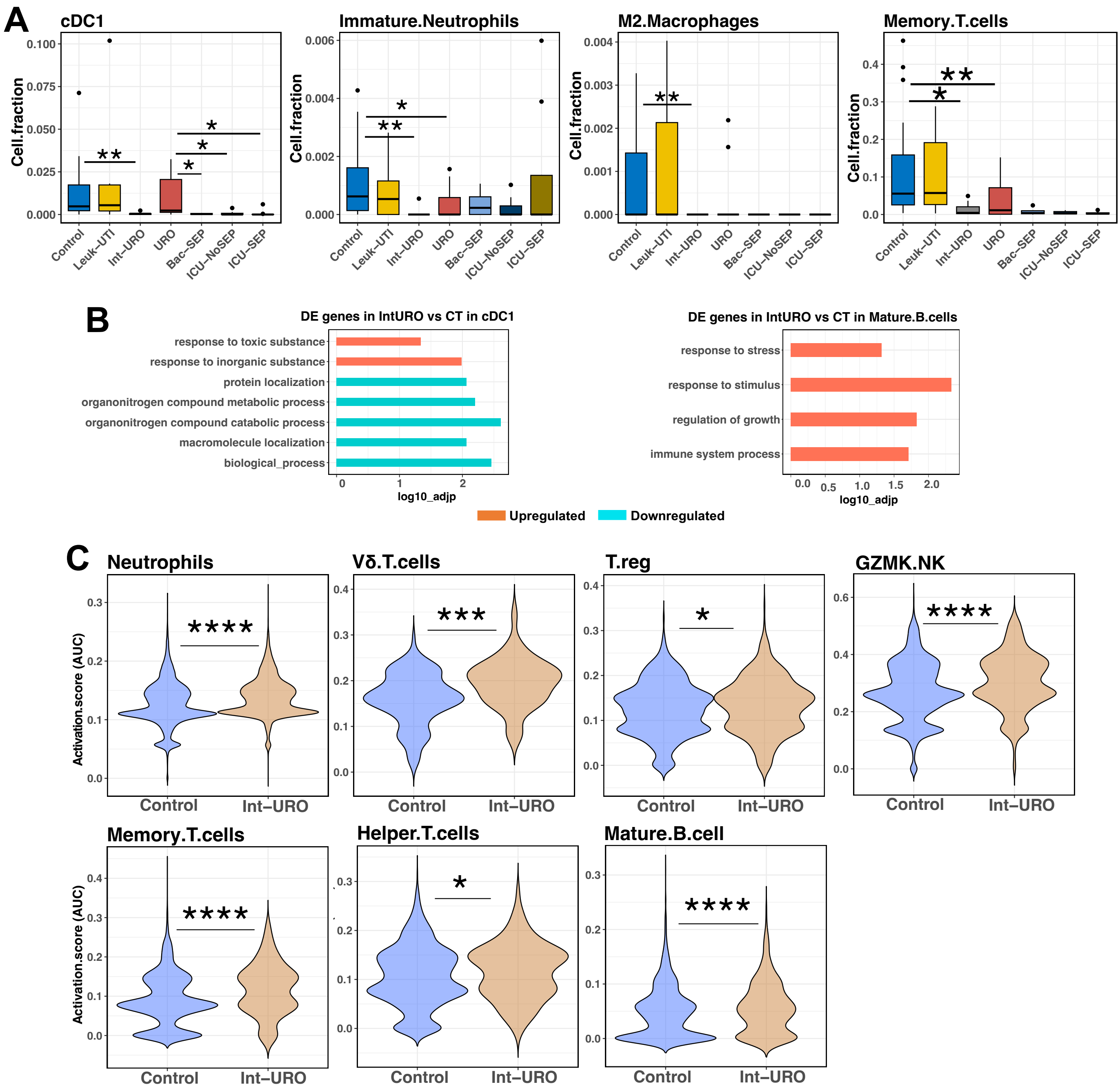

**Figure S10 Cell fraction and immune cell activation activity of immune cells in adult MODS.** (A) Cell fractions of cDC1, immature neutrophils, M2 macrophages, and memory T cells. All these cell types showed decreased abundance in Int-URO compared to control samples. (B) Enriched biological processes identified by upregulated (red) and downregulated (blue) genes in cDC1 and mature B cells when comparing Int-URO with control. Significant enrichment was observed only in these two cell types. (C) Immune cell activation activity in control and Int-URO samples.**** 1e-16< p-value ≤ 1e-05, *** 1e-5 < p-value ≤ 0.0001, ** 0.0001 < p-value ≤ 0.01, and * 0.01< p-value ≤ 0.05.

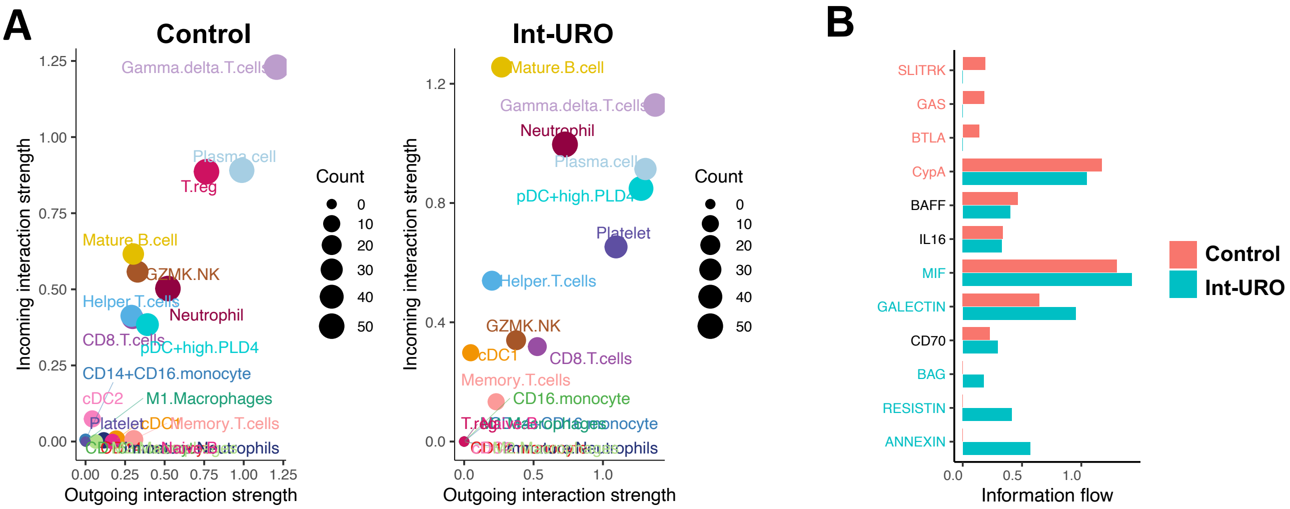

**Figure S11 Pseudotime and cell-cell communications in adult MODS.** (A) Dot plot illustrating the key cell types involved in most cell-cell interactions in control and Int-URO. In control, Vδ T cells were the dominant cell type, whereas in MODS, mature B cells, pDC+highPLD4 cells, and neutrophils were the primary cell types driving cell-cell communication. (B) Bar plot highlighting significant signaling molecules involved in cell-cell communication in control and Int-URO samples.

**Table S1.** Patient demographics and their clinical features.

| Characteristics | Control | MODS |
|---|---|---|
| **Number** | 5 | 5 |
| **Sex** | | |
| Male | 2 (40%) | 2 (40%) |
| Female | 3 (60%) | 3 (60%) |
| **Age** | | |
| Mean (range) | 12.1 (7.5-17) | 11.35 (6-16) |
| **Mortality** | - | 1 |
| **Clinical Features** | | |
| WBC | - | 13.23 (5.3-20.16) |
| Neutrophils | - | 62.1 (23.5-85) |
| Lymphocytes | - | 27.16 (7-70.1) |
| Eosinophils | - | 3.6 (1.1-11) |
| Monocytes | - | 4.66 (0-10.1) |
| Acute Kidney Injury | - | 3 (60%) |
| Respiratory failure | - | 5 (100%) |
| Hepatic Injury | - | 1 (20%) |
| Cardiovascular failure | - | 5 (100%) |
| Shock | - | 5 (100%) |
| LOS | - | 12.68 (3-23) |
